# Sucrose promotes D53 accumulation and tillering in rice

**DOI:** 10.1101/2020.11.10.377549

**Authors:** Suyash B. Patil, Francois F. Barbier, Jinfeng Zhao, Syed Adeel Zafar, Muhammad Uzair, Yinglu Sun, Jingjing Fang, Maria-Dolores Perez-Garcia, Jessica Bertheloot, Soulaiman Sakr, Franziska Fichtner, Tinashe G. Chabikwa, Shoujiang Yuan, Christine A. Beveridge, Xueyong Li

## Abstract

- Shoot branching, a major component of shoot architecture, is regulated by multiple signals. Previous studies have indicated that sucrose may promote branching through suppressing the inhibitory effect of the hormone strigolactone (SL). However, the molecular mechanisms underlying this effect are unknown.
- Here we used molecular and genetic tools to identify the molecular targets underlying the antagonistic interaction between sucrose and SL.
- We showed that sucrose antagonises the suppressive action of SL on tillering in rice and on the degradation of D53, a major target of SL signalling. Sucrose inhibits the expression of *D3*, the orthologue of the arabidopsis F-box protein MAX2 required for SL signalling. Over-expression of *D3* prevents sucrose from inhibiting D53 degradation and enabled the SL inhibition of tillering under high sucrose. Sucrose also prevents SL-induced degradation of D14, the SL receptor involved in D53 degradation. Interestingly, *D14* over-expression enhances D53 protein levels and sucrose-induced tillering.
- Our results show that sucrose inhibits SL perception by targeting key components of SL signalling and, together with previous studies reporting the inhibition of SL synthesis by nitrate and phosphate, demonstrate the central role played by strigolactones in the regulation of plant architecture by nutrients.

## Introduction

Plants are sessile organisms that have evolved the ability to adapt to constantly changing environmental conditions. Shoot branching regulation in angiosperms allows plants to adjust to a given environment, contributes to the overall plant architecture and is considered as an important economic trait for horticulture and agriculture (Wang *et al*., 2018; Li *et al*., 2019; Guo *et al*., 2020). Thus, the study of shoot branching is of major importance for food security, given the increasing global population.

Shoot branching is regulated by different processes including apical dominance, a phenomenon observed in many plants whereby the growing shoot apex inhibits the growth of axillary buds on the same stem. Factors such as herbivory, pruning, and accidental damage to the main shoot break this dominance, allowing lateral buds to grow into branches. Moreover, depending on the environmental conditions, buds can be induced to grow into branches, thus allowing plant architecture to adapt to the prevailing conditions. The shoot tip inhibits axillary bud outgrowth by producing a basipetal flow of auxin in the adjacent stem (Thimann & Bonner, 1933; Domagalska & Leyser, 2011; Barbier *et al*., 2017) and because the rapid growth of the shoot tip diverts nutrients away from axillary buds (Mason *et al*., 2014; Rameau *et al*., 2015; Barbier *et al*., 2019b).

Part of the inhibitory effect of auxin on shoot branching is mediated by the phytohormones, strigolactones (SL). The apically-derived auxin upregulates the expression of strigolactone synthesis genes in the stem (Domagalska & Leyser, 2011; Rameau *et al*., 2015). SL are sensed by DWARF14 (D14), an α/β hydrolase (Arite *et al*., 2009; Hamiaux *et al*., 2012), which then interacts with the F-box protein DWARF3 (D3), the rice orthologue of the arabidopsis MORE AXILLARY GROWTH2 (MAX2), to form a Skp1-Cullin-F-box (SCF) E3 ubiquitin ligase (SCF^D3/MAX2^) complex (Zhao *et al*., 2014). DWARF53 (D53) protein is degraded by the SL-mediated ubiquitination and proteasomal degradation through the D14–SCF^D3^ complex (Jiang *et al*., 2013; Zhou *et al*., 2013). Gain-of-function *d53* mutants in rice and arabidopsis display highly branched phenotypes, and loss-of-function can recover the highly branched phenotype of mutants deficient in SL levels or signalling (Jiang *et al*., 2013; Zhou *et al*., 2013). The TEOSINTE BRANCHED1, CYCLOIDEA, and PCF (TCP) family transcription factor TEOSINTE BRANCHED1 (TB1) acts as a negative regulator of tillering and is known as an important hub, integrating different signals, including SL, that induce *TB1* gene expression (Takeda *et al*., 2003; Aguilar-Martínez *et al*., 2007; Minakuchi *et al*., 2010; Dun *et al*., 2012; Wang *et al*., 2019a).

Auxin treatment is not always enough to restore apical dominance after shoot tip removal (Cline, 1996; Beveridge, 2000; Morris *et al*., 2005; Barbier *et al*., 2021). As demonstrated upon decapitation in pea, auxin depletion in the stem does not correlate with initial bud growth (Morris *et al*., 2005; Ferguson & Beveridge, 2009). In this species, this initial bud outgrowth has been correlated with rapid remobilisation of carbohydrates towards the buds (Mason *et al*., 2014; Fichtner *et al*., 2017). Beyond their trophic role, sugars also act as signalling molecules, allowing plants to adjust their metabolism, growth and development to their environment (Rolland *et al*., 2006; Sakr *et al*., 2018; Fichtner *et al*., 2021b). A signalling role for sugars in bud outgrowth and shoot branching has been reported for different species (Takahashi *et al*., 2014; Barbier *et al*., 2015b,a). The low abundant metabolite trehalose 6-phosphate (Tre6P), a sucrose-specific signalling molecule (Fichtner & Lunn, 2021), accumulates rapidly in pea buds upon decapitation (Fichtner *et al*., 2017) and promotes shoot branching in arabidopsis (Fichtner *et al*., 2021a). The HEXOKINASE1 (HXK1) signalling pathway (Moore *et al*., 2003) was also recently shown to promote shoot branching in arabidopsis and to interact with both, the CK and SL pathways (Barbier *et al*., 2021). A recent study comparing transcriptomes of annual and perennial plants correlated the expression of genes involved in carbon starvation with bud dormancy (Tarancón *et al*., 2017). In different eudicots, sugar supply to the plant can promote shoot branching (Mason *et al*., 2014; Barbier *et al*., 2015b; Dierck *et al*., 2016); however, this has not yet been reported for monocot plants.

The interactions among sugars and hormones during the control of bud outgrowth are not yet fully resolved. Recent studies have indicated that sucrose can antagonise the effect of auxin by inhibiting SL perception to promote bud outgrowth (Bertheloot *et al*., 2020) and that the promotion of growth by cytokinins may only be effective under conditions where sugars are not readily available (Barbier *et al*., 2015b; Salam *et al*., 2021). Interactions between sugars and strigolactones have been recently highlighted in rice during the control of shoot architecture by the circadian clock (Wang *et al*., 2020). Interestingly, sucrose application to single-node cuttings of rose buds can suppress *MAX2* and *BRANCHED1* (*BRC1*, the arabidopsis orthologue of *TB1*) gene expression (Barbier *et al*., 2015b; Wang *et al*., 2019b). In sorghum (*Sorghum bicolor*), defoliation and shade treatments, which decrease sugar availability, inhibit bud outgrowth and up-regulate *MAX2* expression (Kebrom *et al*., 2010; Kebrom & Mullet, 2015).

The aim of this study was to test whether sugar availability affects SL-induced tillering inhibition in rice and to make the first steps towards identifying the molecular components involved. Using physiological experiments and genetic tools, we sought to identify which components of SL signalling are targeted by sucrose during tillering in rice, and bud outgrowth in pea. Since both sugar and SL levels in plants are controlled by environmental factors, this study will shed light on how environmental factors may regulate branching and tillering at the molecular level.

## Materials and Methods

### Plant material and growth conditions

For rice, available lines created in different backgrounds were used as indicated in the figure legends, and the corresponding wild types were used as controls. The tiller development assay in response to sucrose and GR24 was performed using the *japonica* cultivar HuaiDao5(Patil *et al*., 2019). The seeds were sterilised as per the method described earlier (Zhao *et al*., 2014) with slight modifications. In brief, de-husked seeds were sterilised with 30% NaClO solution in a shaker for 30 minutes and then washed with sterilised de-ionised water at least five times. The seeds were directly sown on the solidified (0.5% agar) half-strength MS media with different sucrose concentrations with adjusted pH of 5.8. For the SL treatments, rac-GR24 was used in all the experiments (CX23880, Chiralix). The plants were grown on the different sucrose concentrations for three weeks with or without 1 μM GR24 under a 16-hr light (200 μmol m^-2^ s^-1^)/8 hr dark cycle at 28°C in a growth chamber. The GR24 and the corresponding treatment combinations were replaced at weekly intervals, maintaining strictly sterile conditions.

The calli of wild-type (WT) plants (HuaiDao5) grown on NB media plates at 28°C in the dark were used for the D53 protein degradation assay. The *D3* and *D14* over-expressing lines and their corresponding mutant and WT lines were used from the earlier work (Zhao *et al*., 2014). The lines were maintained in field condition at the experimental station of Shandong Rice Research Institute, Shandong, China. The lines used for sucrose sensitivity assay consisted of *d3* (*s2-215*^*Q393Stop*^) (Patil *et al*., 2019) and *d14* (*htd-2*) (Liu *et al*., 2009) mutants in the Nipponbare WT background.

For pea, decapitation and sucrose petiole feeding experiments were performed on the Torsdag L107 background. *In vitro* sucrose treatment with single nodes was performed as described earlier (Bertheloot *et al*., 2020) in the *rms3, rms4* mutants and their corresponding WT Terese. For the decapitation experiment, plants were grown in a glasshouse with a controlled environment (Fichtner *et al*., 2017). For the sucrose petiole feeding experiment, plants were grown in a growth chamber with 16 hrs of light (125 μmol m^-2^ s^-1^) at a temperature of 22°C during the day and of 20°C at night.

### Rice callus induction

Husks were removed from rice seeds which were then sterilized with the 30% NaClO solution and washed for multiple times with sterilized deionized water. The seeds were then placed on solidified NB media petri-plates in the dark at 28°C for one week. The embryonic calli were then separated and multiplied on fresh NB media plates maintained at 28°C in the dark. To prepare 1 L NB medium, 4.1 g NB basal medium (Phytotech lab), 2 g Casein hydrolysate, 4 g L-Proline, 2 g L-Glutamine, 200 µl 2-4-Dichlorophenoxyacetic acid (10 mg ml^-1^), 30 g sucrose (adjusted depending on experimental needs) and 3 g Phytagel were mixed together (pH = 5.8).

### Protein quantification assay using callus tissues and tiller buds

The calli were grown on NB media plates containing different sucrose concentrations. Around 250 mg callus tissue was ground in liquid nitrogen, using a mortar and pestle to make a fine powder. The powder was transferred to 1.5 ml microcentrifuge tubes and mixed with 250 µl of ice-cold TBT buffer (100 mM AcOK, 20 mM KHEPES pH 7.4, 2 mM MgCl2, 0.1% Tween-20, 1 mM DTT, and 0.1% protease inhibitor cocktail). The antibody preparation, samples preparation, and protein blots were developed as per the method described earlier (Jiang *et al*., 2013). The D53-specific polyclonal antibody produced in mouse was used for immune detection following 1:1000 dilution in non-fat dairy milk. Anti-HSP82 or anti-actin was used as a loading control following 1:3000 dilution. Horse-radish peroxidase-conjugated anti-mouse lgG was used as a secondary antibody (CWBIO, Beijing, China) following 1:3000 dilution. The western blots were developed using Tanon™ High-sig ECL western blotting Substrate (Cat. no:180-5001) with a Tanon 6100 chemiluminescent imaging system. For each Western blot, the band intensity of the target and loading control proteins was determined using the “Gels” analysing tool in ImageJ. The values of the target proteins were then normalized by the values of the loading control, and the values were presented relative to the control condition for each genotype to ease comparison across treatments.

### Gene expression analysis

For rice, shoot base tissues (0.5 cm) were harvested from two-week-old seedlings grown hydroponically. Total RNA was isolated using RNAprep Pure Plant Kit (Tiangen, Beijing, China, cat. no. DP432) following the manufacturer’s instructions. Reverse transcription of 500 ng RNA was performed using the Vazyme, HiScript II Q Select RT SuperMix for qPCR (cat no. R233). Real-time quantitative PCR was performed by Vazymes Cham Q QPCR reagent kit (cat no. Q331) using the ABI Prism 7500 Sequence Detection System as per the program recommended by both the instrument and the reagent company. Transcript levels were detected by CT values relative to *ACTIN1* as a reference gene. All the primer sequences used in this study are listed in Supplementary Table S1. Gene expression in pea buds was monitored using a phenol/chloroform free CTAB-based method as described earlier (Barbier *et al*., 2019a).

### *Nicotiana benthamiana* leaf agroinfiltration and *in vitro* luciferase activity assay

The pCambia1200 vector was modified by integrating Firefly luciferase and Renilla luciferase coding sequences driven by the CaMV35S promoter in addition to the hygromycin resistance marker. The vector then called pCambia1200 35S-LUC had both transient and stable expression capabilities (Figure S2) (Sun *et al*., 2021). The coding sequence of the *D53* gene was amplified using D53 LUC-F-GGGCGGAAAGGAATTCATGCCCACTCCGGTGG and D53 LUC-R-TAGATCCGGTGGATCCTCAACAATCTAGAATTATTCTTGGCGGGAG primer pairs and cloned at the C-terminal region of the Firefly luciferase gene using *Eco*RI and *Bam*HI restriction sites following the In-fusion**®** cloning system by Clonetech, Takara biotech, Japan. The plasmids were transformed into the Agrobacterium strain EHA105 to transfect into *Nicotiana benthamiana* leaves for the transient expression of the D53-Firefly luciferase fused protein. This vector also consists of a Renilla luciferase gene as an internal control which was used to quantify the relative amount of the D53 protein levels. The Agrobacterium strain was transfected into the *N. benthamiana* leaves as per the methods described earlier (Chen *et al*., 2008). To check the effects of sucrose and GR24 on the transiently expressed D53 levels, different treatment combinations were infiltrated directly into the leaves already transfected with the Agrobacterium strain harbouring the luciferase-fused D53 construct. The leaves were then harvested in liquid nitrogen, and the activities of Firefly luciferase and Renilla luciferase were determined using the Dual-luciferase® reporter assay system from Promega (cat. no. E1910). The LUC activity was calculated by normalising Firefly luciferase values with Renilla luciferase and was presented as relative luciferase values.

## Results

### Sucrose and strigolactones interact antagonistically to regulate tillering in rice

In dicotyledonous models, sucrose has been shown to promote axillary bud outgrowth and alleviate the inhibitory effect of SL during this process (Dierck *et al*., 2016; Bertheloot *et al*., 2020). However, this antagonistic interaction has not been reported in monocotyledons plants, which have a different architecture compared with eudicots, and are evolutionarily distant. To evaluate sucrose and SL effects on tillering bud outgrowth in rice, we grew wild-type (WT) Huaidao-5 plants hydroponically with different sucrose concentrations with or without 1 μM rac-GR24 (a synthetic SL analogue). To be in a physiological range of sucrose concentrations, we added up to 4% sucrose (∼120 mM), which is lower than the endogenous sucrose levels reported in rice (between 200 mM and 600 mM) (Hayashi & Chino, 1990). We observed that sucrose triggered bud elongation in a dose-dependent manner with or without addition of rac-GR24 (Figure 1). In the absence of sucrose, tiller bud elongation remained suppressed. With 2% sucrose, GR24 strongly inhibited tiller bud elongation (73% inhibition) (Figure 1B). However, with 4% sucrose, the inhibitory effect of GR24 on tiller bud elongation was reduced to 30% inhibition. A two-way ANOVA demonstrated that the effect of GR24 largely depends on the concentration of sucrose (*p*-value = 0.000019). In contrast to sucrose, sorbitol only had a very small promoting effect on tillering and barely alleviated the inhibitory effect of GR24, showing that the effect of sucrose was largely independent of an osmotic effect (Supplementary Figure S1). These phenotypic data support a similar hypothesis as proposed previously for selected eudicots, namely that sucrose and SL interact antagonistically to regulate tillering.

**Figure 1.**
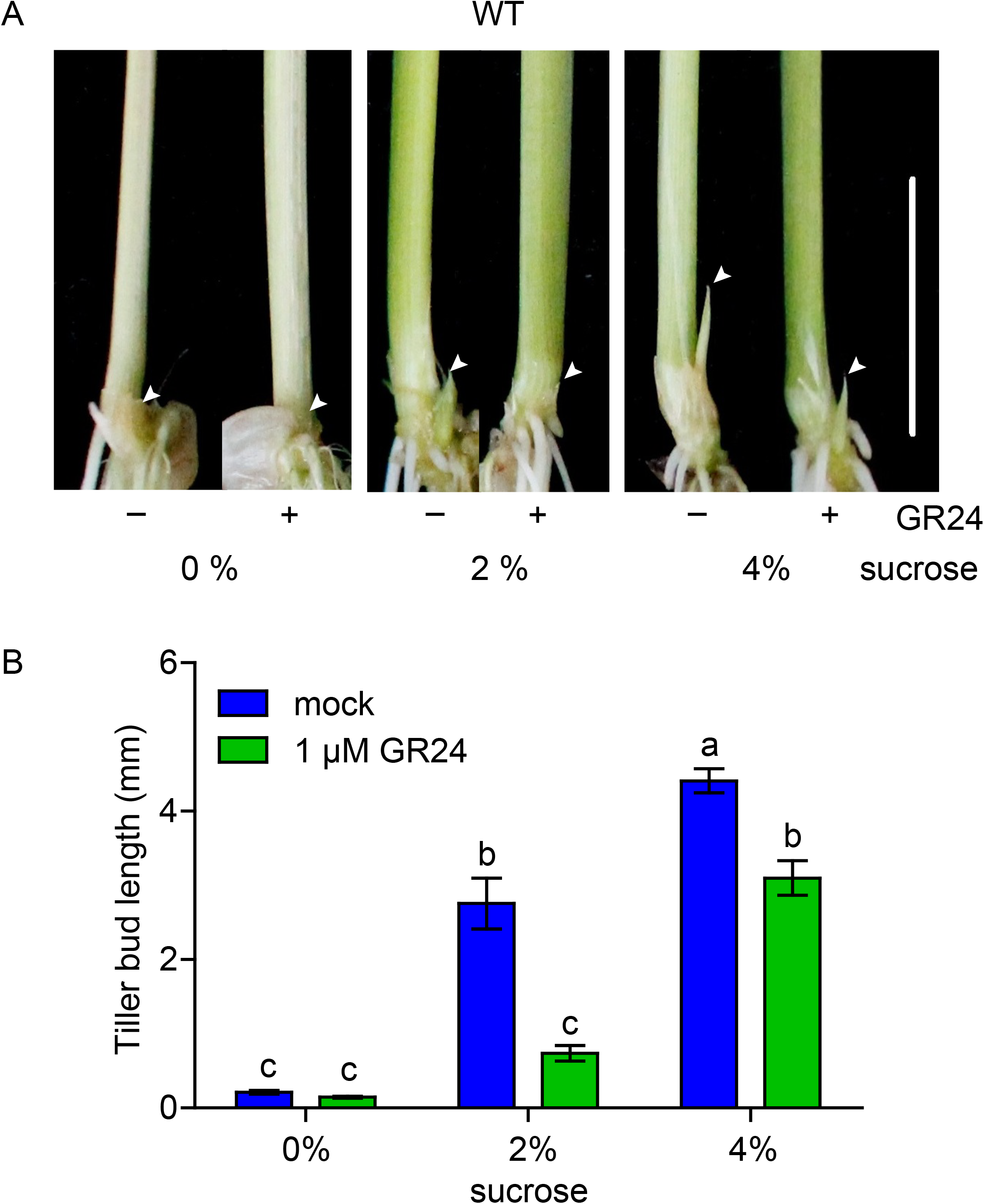
Sucrose alleviates the inhibitory effect of GR24 on tiller bud outgrowth in rice. (**A**) Representative tiller buds and (**B**) length of tiller buds of the WT (Huaidao 5) grown under different sucrose concentrations with or without 1 μM GR24 for 3 weeks. Scale bar represents 10 mm. Different lower-case letters denote significant differences (p<0.05, one-way ANOVA following Tukey’s test for multiple comparisons). Error bars represent ± SE (n > 8).

### Sucrose inhibits GR24-induced degradation of D53 protein in rice

The D53 protein and its orthologues in arabidopsis SMXL6, 7 and 8 play a crucial role in SL-mediated shoot branching in rice (Jiang *et al*., 2013; Zhou *et al*., 2013) and arabidopsis (Soundappan *et al*., 2015; Wang *et al*., 2015), respectively. Since sucrose reduces the SL response, we proposed that sucrose might promote D53 accumulation. We therefore tested the impact of sucrose on D53 accumulation. Since dormant tiller buds are very small and D53 protein levels are difficult to detect in shoot tissues, we first used rice calli, which have previously been successfully used for this purpose (Jiang *et al*., 2013). In the absence of GR24 in the growth medium, the D53 protein levels strongly increased with the increasing sucrose concentration (Figure 2A)

**Figure 2.**
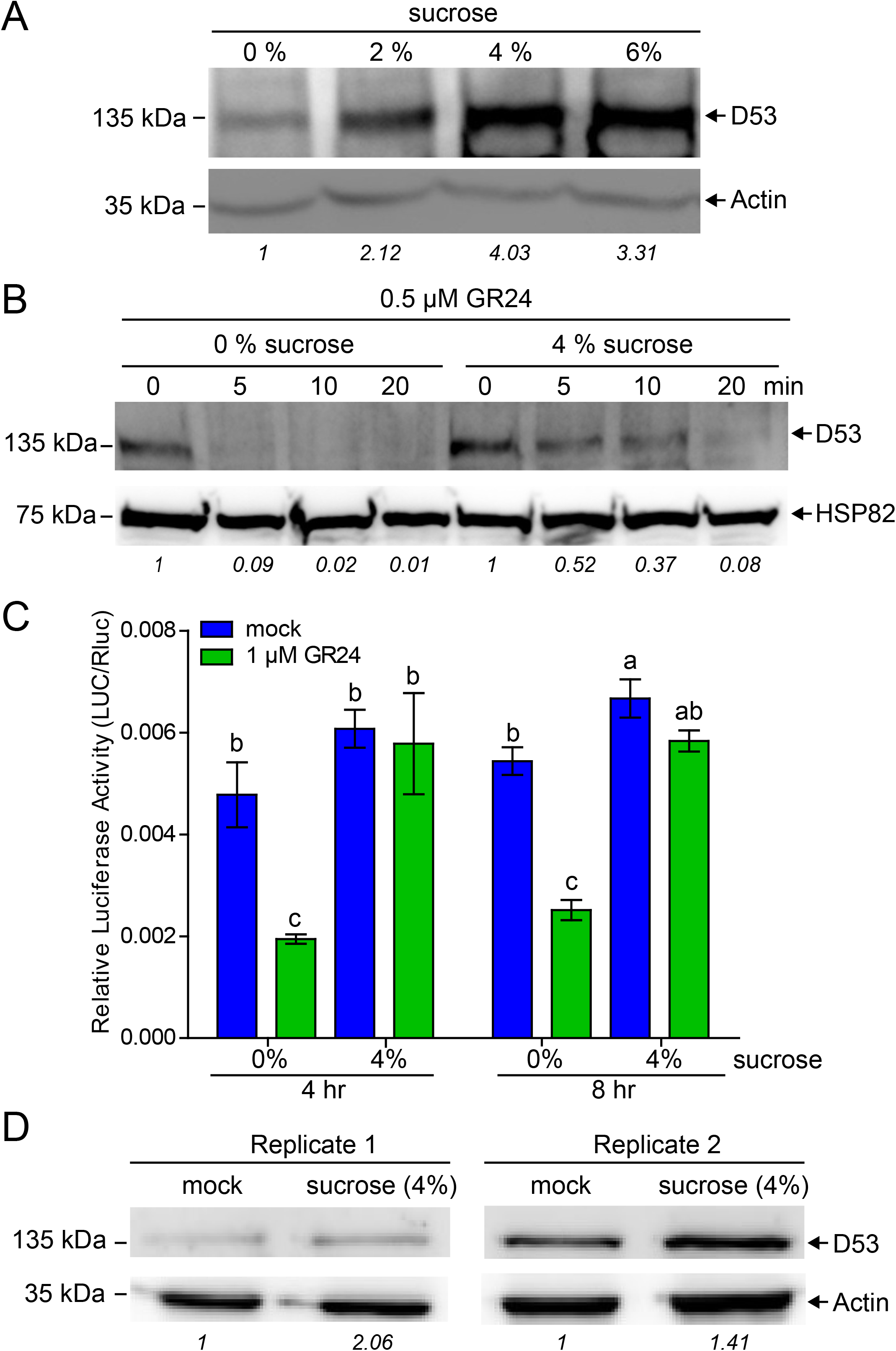
Sucrose alleviates the GR24-induced D53 degradation. (**A**) Western blot showing D53 protein accumulation in rice calli grown for 4 weeks on different sucrose concentrations. (**B**) D53 protein degradation in the WT rice calli initially grown on NB plates containing 4% sucrose and later shifted to liquid media with or without 4% sucrose containing 0.5 μM GR24 for different time points. (**C**) Transiently expressed luciferase-D53 protein in *Nicotiana benthamiana* leaves subjected to 1 μM GR24 treatment with or without 4% sucrose for different time points. Luciferase readings were normalised with Renilla luciferase readings. Values are mean ± SE (n = 4). Different lower-case letters denote significant differences (p<0.05, one-way ANOVA following Tukey’s test for multiple comparisons) (**D**) Western blot showing D53 accumulation in isolated tillers buds (< 3 mm length) of 3-month-old rice (Nipponbare) plants treated with or without 4 % sucrose for 1 hour. Numbers in italics below the blots indicate the relative band intensity of D53 normalized by the intensity of Actin or HSP82 bands.

We then tested the effect of sucrose on D53 in the presence of GR24. As the calli grown on different sucrose concentrations accumulate different levels of D53, calli grown on 4% sucrose plates showing a similar amount of D53 protein were used. The calli were washed twice with sterile water to remove the exogenous sucrose before being transferred to liquid media containing either no sucrose, or 4% sucrose. After 30 minutes of stabilisation, 0.5 μM GR24 was supplemented to the 0% and 4% sucrose treatments. In the absence of sucrose, D53 protein was degraded within 5 minutes of treatment with GR24. However, in the presence of 4% sucrose, it took 20 minutes for GR24 to lead to a similar degradation of the D53 protein (Figure 2B).

To confirm this result, we tested OsD53 degradation in response to sucrose and GR24 in *Nicotiana benthamiana* leaves transiently expressing an *OsD53* coding sequence fused to a LUCIFERASE (LUC) reporter (Supplementary Figure S2). D53 protein accumulation was assessed by measuring the LUC activity normalised with Renilla luciferase values. Without GR24, sucrose had a minor effect on LUC activity which was only significantly enhanced by sucrose at 8 hrs. In the absence of sucrose, the LUC activity was lower in the presence rather than in the absence of GR24 at 4 hr and 8 hr after hormone or control treatment. However, in the presence of 4% sucrose, the LUC activity was similar with or without GR24 (Figure 2C). Again, these observations show that D53 protein levels are maintained at higher levels in the presence of sucrose and that the degradation rate of D53 in response to GR24 is lower under these sucrose conditions.

To observe whether results from calli and agroinfiltrated *N. benthamiana* may be relevant *in planta*, we measured D53 proteins levels in dormant tiller buds (length < 3 mm) harvested from plants fed hydroponically with 4% sucrose for one hour, and in which tiller buds were only subjected to the plant’s endogenous SL levels. The result indicated that supplying 4% sucrose to rice plants grown hydroponically promoted the accumulation of the D53 protein in dormant tiller buds (Figure 2D), further supporting the hypothesis that sucrose promotes D53 accumulation during tillering regulation.

### Sucrose inhibits the expression of genes involved in strigolactone signalling

D53 promotes the outgrowth of tillers by inhibiting the expression of the TCP transcription factor gene *TB1* (Takeda *et al*., 2003; Minakuchi *et al*., 2010). Given the accumulation of D53 proteins in response to sucrose (Figure 2), we predicted that sucrose treatment should suppress *TB1* expression. To test this, we measured *TB1* expression in rice calli treated with a range of sucrose concentrations. The results show that sucrose inhibited the expression of *TB1* in rice callus tissues in a dose-dependent manner (Figure 3A).

**Figure 3.**
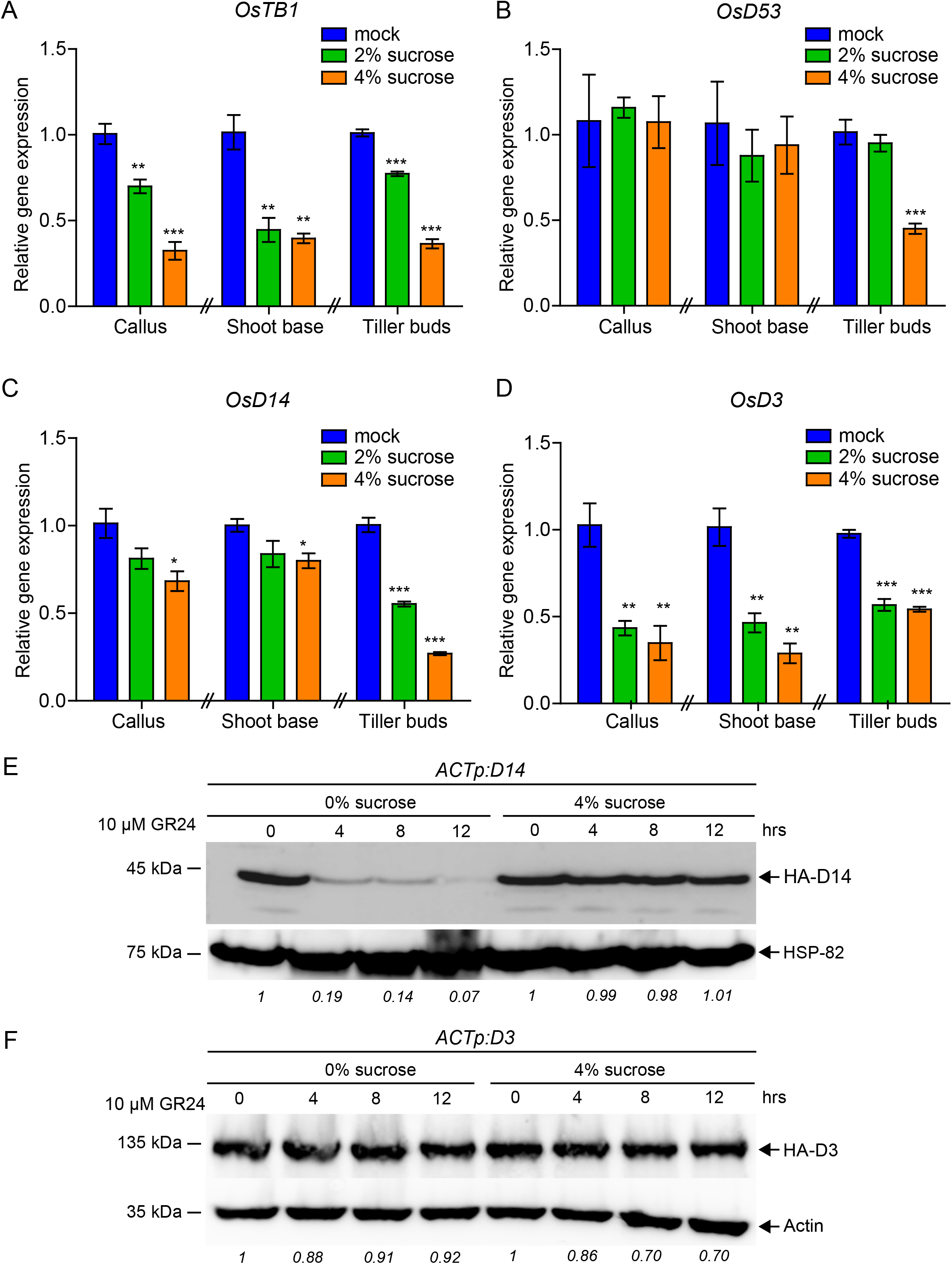
Sucrose down-regulates key genes in the SL signalling pathway. (**A**) Effect of different sucrose concentrations on the expression of *TB1*, (**B**) *D53*, (**C**) *D14* and (**D**) *D3* in the callus (grown for three weeks), shoot base tissues (three weeks old plants), and isolated tiller buds (<3 mm length) of two-month-old rice plants grown hydroponically with different sucrose concentrations for 24 hrs. Values are mean ± SE (n = 3). Each replicate consists of 8 biologically independent samples. Significant levels: * p < 0.05, ** p < 0.01; indicated by Student’s *t*-Test. (**E**) Effect of sucrose on GR24-mediated degradation of HA-tagged D14 fused protein levels detected by immunoblotting with an anti-HA tag monoclonal antibody. (**F**) Effect of sucrose on GR24-mediated degradation of HA-tagged D3 fused protein levels detected by immunoblotting with an anti-HA tag monoclonal antibody. Numbers in italics below the blots indicate the relative band intensity of D14 or D3 normalized by the intensity of Actin or HSP82 bands.

We then determined whether the change in D53 protein levels could be explained by a change in *D53* expression at the transcript level. Our results indicate that, in contrast to *TB1* expression and the D53 protein level, *D53* gene expression is not responsive to sucrose (Figure 3B). These observations suggest that sucrose promotes D53 protein accumulation through a post-transcriptional mechanism.

We then tested whether the observations made on calli were relevant to tissues where the regulation of tillering occurs. To do so, we harvested dormant tiller buds (length < 3 mm) and shoot bases (length = 5 mm), enriched in buds and stem. As in calli, sucrose treatment through hydroponic media down-regulated the expression of *TB1* (Figure 3A) in tiller buds and in shoot bases. The expression of *D53* was not affected by sucrose in calli or in the shoot base, but was repressed by 4% sucrose in tiller buds. This is consistent with the negative correlation reported for D53 protein level and *D53* gene expression (Zhou *et al*., 2013).

Given that sucrose reduces SL response in buds (Dierck *et al*., 2016; Bertheloot *et al*., 2020) (Figure 1) and enhances the level of D53 protein (Figure 2), we predicted that sucrose may affect components of the SCF complex formed by D14 and D3, which are required for D53 protein degradation (Zhou *et al*., 2013; Zhao *et al*., 2014). Consistent with such a role of D3 and D14 in sucrose regulation of D53, *D3* and *D14* gene expression was significantly reduced by sucrose in calli, tiller buds and shoot bases, although the inhibition of *D14* in shoot bases was milder compared to the inhibition of *D3* expression in this tissue (Figure 3C,D).

We then tested whether sucrose could also directly regulate D14 or D3 protein levels. To do so, we tested the impact of 4% sucrose on D14 and D3 accumulation in calli of transgenic lines over-expressing HA-tagged *D14* or *D3* driven by the constitutively active *OsACTIN1* promoter (*ACTp*) in the *d14* and *d3* mutant background, respectively (Supplementary Figure S3). In these lines, we expected stable synthesis of the tagged proteins. To establish this approach, we first observed that the HA-tagged over-expressing lines complemented the tiller number and almost fully complemented the plant height phenotypes of the corresponding *d14* and *d3* mutants (Supplementary Figure S4). In the absence of sucrose, GR24 led to the almost complete degradation of the D14 protein over 12 hrs in the HA-D14 over-expression line (Figure 3E). This indicates that the GR24-induced degradation rate of the HA-D14 protein in the absence of sucrose exceeds the stable rate of HA-D14 synthesis in this line. Strikingly, in the presence of sucrose, GR24 did not affect the levels of HA-D14 (Figure 3E). This indicates that sucrose antagonises GR24-induced degradation of D14. In contrast, D3 protein levels in the HA-D3 over-expression line were quite stable in response to GR24 and sucrose (Figure 3F). Altogether, this suggests that sucrose inhibits the GR24-induced D14 degradation and does not directly regulate D3 protein levels.

### D3/OsMAX2 over-expression prevents the regulation of D53 by sucrose

If sucrose acts via D14 and/or D3 to regulate D53 protein accumulation as indicated above (Figure 3A-D), we would predict that over-expression of one or both of these two genes would prevent sucrose from promoting D53 accumulation and may prevent sucrose-induced tillering. To test the sucrose response over a short time frame, we grew the calli of *ACTp*:*D3, ACTp*:*D14* over-expression lines on NB medium plates supplemented with 1% sucrose. Since these lines have been created in different backgrounds (GSOR300002 and GSOR300192, respectively), these cultivars were used as controls. The calli were then washed and rinsed for 60 min with sterile water. The calli were then shifted to liquid media containing 4% sucrose and collected after 0, 2, 4 and 8 hrs to determine D53 protein levels. In two WT backgrounds, an increase in D53 levels was observed in response to sucrose (Figure 4A,B). Constitutive over-expression of *D14* or *D3* diminishes the response of D53 to sucrose but in two different and opposite ways. *D14* overexpression causes a high level of D53 protein, regardless of the sucrose supply. In contrast *D3* overexpression prevents accumulation of D53 under sucrose treatment. These results indicate that *D3* over-expression prevents sucrose-induced D53 accumulation, while *D14* over-expression mimics the effect of sucrose supply on D53 protein levels.

**Figure 4.**
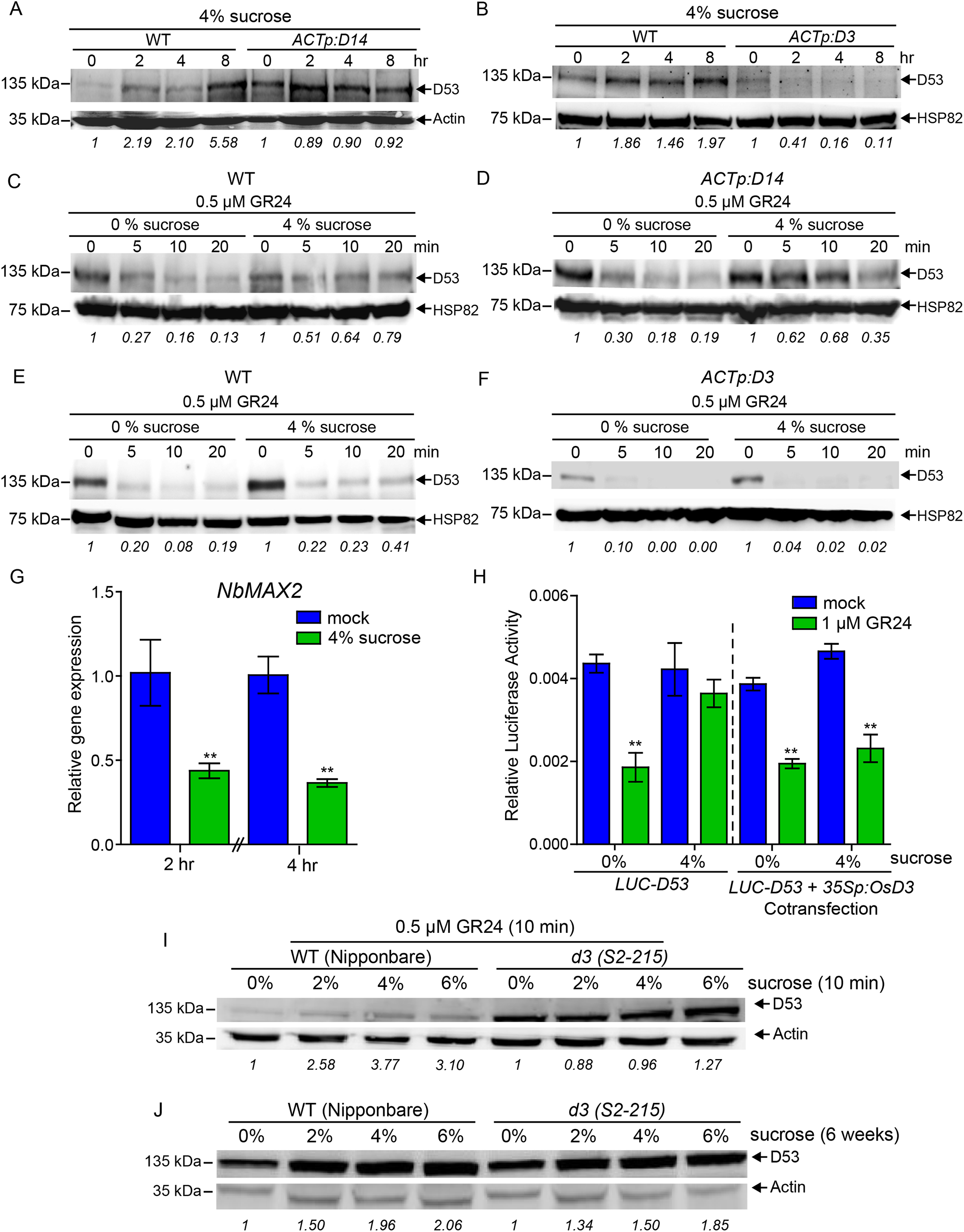
Over-expression of *D3*, but not *D14*, leads to D53 degradation in the presence of sucrose. (**A**) Effect of sucrose (4%) on D53 accumulation at different time points in WT and *D14* over-expressing line, and (**B**) WT and *D3* over-expressing line detected by immunoblotting with an anti-D53 polyclonal antibody. (**C**) Effect of sucrose on D53 degradation in the calli of WT (GSOR300192), (**D**) *D14* over-expressing line, (**E**) WT (GSOR300002), and (**F**) *D3* over-expressing line initially grown on NB plates containing 4% sucrose and later shifted to liquid media with or without 4% sucrose containing 0.5 μM GR24 for different time points detected by immunoblotting with an anti-D53 polyclonal antibody. (**G**) Effect of sucrose on *Nicotiana benthamiana MAX2* (*NbMAX2*) expression in *Nicotiana benthamiana* leaves infiltrated with or without sucrose solution (4%) at different time points. Values are mean ± SE (n=3). (**H**) Transiently expressed luciferase-D53 protein alone and co-transfected with 35S-OsD3 protein in *N. benthamiana* leaves subjected to 1 μM GR24 treatment with or without 4% sucrose for different time points. Luciferase readings were normalised with renilla luciferase readings. Values are mean ± SE (n = 3). Significant levels: **p < 0.01; indicated by Student’s *t*-Test. (**I**) D53 accumulation in rice in WT and *d3* mutant calli grown NB plates containing 4% sucrose and later shifted to liquid media with 0.5 μM GR24 and a range of sucrose concentrations for 10 min. (**J**) D53 accumulation in rice in WT and *d3* mutant calli continuously grown on different sucrose concentrations for 6 weeks without exogenous GR24. Numbers in italics below the blots indicate the relative band intensity of D53 normalized by the intensity of Actin or HSP82 bands.

Since sucrose and SL have an antagonistic effect on D53 protein levels (Figure 2B-C), we measured D53 protein levels in rice calli over-expressing *D14* and *D3* in response to both GR24 and sucrose. As observed in Figure 2, the effect of GR24 on D53 degradation was delayed by sucrose treatment in the WTs (Figure 4C and 4E). *D14* overexpression did not prevent the promoting effect of sucrose on D53 levels (Figure 4D). In contrast, *D3* over-expression led to total degradation of D53 after 5 minutes of GR24 treatment, regardless of the sucrose concentration in the medium (Figure 4F). These results suggest that *D3* over-expression, but not that of *D14*, prevents sucrose from antagonising the SL-induced D53 degradation.

Similar findings to those obtained in rice calli were also observed in *N. benthamiana* leaves. In this system the native *N. benthamiana MAX2/D3* expression was also inhibited by sucrose (Figure 4G). We thus tested whether over-expressing *D3* in *N. benthamiana* leaves would also prevent sucrose from antagonising the effect of SL on D53 accumulation. We therefore followed the same procedure as described in Figure 2C and co-transfected D53-LUC with a *35Sp:D3* construct into *N. benthamiana* leaves. In this system, 4% sucrose almost totally alleviated the effect of GR24 on D53 degradation when D53-LUC was solely transfected (Figure 4H). However, when the LUC-D53 construct was co-transfected with the *35Sp:D3* over-expressing construct, sucrose could not prevent the negative effect of GR24 on D53 levels (Figure 4H). The same trend was also observed in a second independent experiment (Supplementary Figure S5). These results further support the hypothesis that sucrose alleviates the inhibitory effect of GR24 on D53 protein levels by inhibiting *D3/MAX2* expression.

We then tested whether sucrose could still have a promoting effect on D53 levels in the *d3* mutant. To do so, we grew WT and *d3* rice calli on 4% sucrose for three weeks, rinsed them with water and transferred them on a range of sucrose concentrations with 0.5 μM GR24 for 10 min. The result shows that D53 accumulates a lot more in the *d3* than in the WT background, as expected in presence of GR24 (Figure 4I). However, we could observe a strong accumulation of D53 in response to sucrose in the WT calli, while no obvious pattern was observed in the *d3* background. This suggests that the positive effect of sucrose on D53 accumulation is visible only if D3 is functional, at least in presence of SL and on this short time-frame.

To further investigate whether sucrose could act through D3-independent pathways to regulate D53 protein levels, we measured D53 in WT and *d3* calli grown on a range of sucrose for six weeks without SL (Figure 4J). In these conditions we could observe a positive effect of sucrose in both backgrounds. We also measured *TB1* expression in WT, *d3* and *d14* calli fed for 24 hrs with a range of sucrose concentrations without SL (Supplementary Figure S6). The results indicate that *TB1* expression remains responsive to sucrose in the three backgrounds. Altogether, these results suggest that sucrose may also act through D3-independent pathway(s) to regulate D53 protein levels and *TB1* expression in calli, at least without exogenous SL supplementation.

### *D3/OsMAX2* over-expression prevents the promoting effect of sucrose on tillering

Given that D53 promotes tillering (Jiang *et al*., 2013; Zhou *et al*., 2013) and that over-expression of *D3*, but not *D14*, prevents the sucrose-induced accumulation of D53 (Figure 4), we tested whether *D3* overexpression in particular would prevent sucrose-induced tillering. WTs and lines over-expressing *D3* and *D14* were grown on 0%, 0.5%, 2% and 4% sucrose media with or without 1 μM GR24 for three weeks. As recorded in our previous experiment (Figure 1), 4% sucrose could antagonise the inhibitory effect of GR24 on tiller bud elongation in the WT lines (Figure 5 A,C). As observed in Figure 1B, two-way ANOVA comparison demonstrated that the impact of GR24 significantly depends on sucrose in the two WT backgrounds (*p*-value = 0.00004 for GSOR300192 and *p*-value = 0.0348 for GSOR300002). This antagonistic effect between GR24 and sucrose was also observed in the *D14* over-expression line (Figure 5B). Strikingly, *D14* over-expression strongly increased the effect of sucrose on tiller length (Figure 5 A,B). In contrast with WT and the *D14* over-expression line, *D3* over-expression inhibited the bud response to sucrose most prominently in presence of GR24 (77% inhibition with 4% sucrose) (Figure 5D). Multiple-way ANOVA comparison revealed that the interaction between sucrose and GR24 did not depend on D14-overexpression (*p*-value = 0.33), whereas it showed that this interaction is highly dependent on D3 over-expression *p*-value = 0.000002). Our data indicate that *D3* over-expression inhibits tiller bud elongation in the absence of sucrose, contrasting with results observed in arabidopisis (Stirnberg *et al*., 2007). The difference may be due to the fact that, in the study in arabidopsis, the branching phenotype was recorded as the number of branches at the end of the plant’s life, whereas our study in rice captured branching at the beginning of the tiller development. Altogether, these results indicate that *D3* over-expression prevents sucrose from alleviating the inhibitory effect of SL on tillering and that *D14* over-expression promotes sucrose-induced tiller bud elongation.

**Figure 5.**
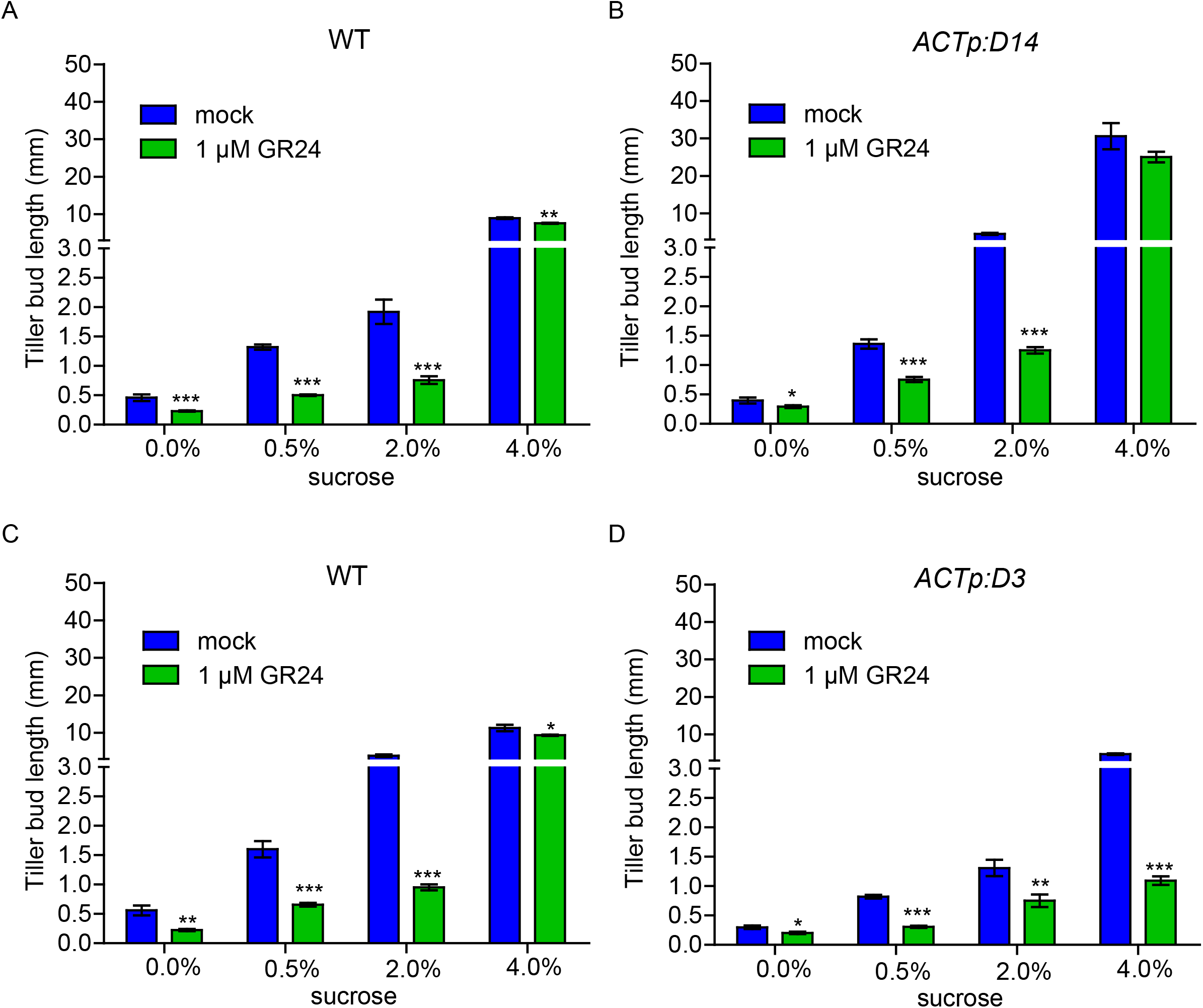
*D3* over-expression prevents sucrose from inhibiting the SL-induced tillering suppression. (**A**) Effect of sucrose on tiller bud elongation in the WT (GSOR300192), (**B**) *D14* over-expression line, (**C**) WT (GSOR300002) and (**D**) *D3* over-expressing line grown with or without 1 μM GR24 for 3 weeks. Values are mean ± SE (n = 10). Significant levels: ***p < 0.001; **p < 0.01, *p < 0.05; indicated by Student’s *t*-Test.

### Strigolactone signalling mutants are less sensitive to low sucrose concentrations and remain responsive to sucrose

Our results suggest that the inhibition of tillering in response to low sucrose is due to high strigolactone signalling. If it holds true, the strigolactone signalling mutant *d14* and *d3* should be less inhibited by low sucrose concentrations. We therefore tested this by measuring the response of the loss-of-function *d3* and *d14* mutants (Supplementary Figure S7) to different sucrose concentrations supplied hydroponically (Figure 6). The buds of the *d3* mutant responded more than the WT to low sucrose concentrations including 0% exogenous sucrose (Figure 6A,C,D). The buds of *d14* mutant also grew better on 0% sucrose than the WT. However, the length of *d14* tiller buds were much smaller than the buds of *d3*, and also showed enhanced growth on the difference sucrose concentrations, but this effect was not always significantly different from the WT and was always lower than *d3*. We could also observe that the buds of *d14* and *d3* mutants were still responsive to the increase in sucrose concentration. This observation is in line with our results showing that sucrose could promote D53 accumulation in a D3-independent manner in absence of exogeneous SL (Figure 4J). Altogether, these data indicate that disrupting SL signalling components, especially D3, decreases the sensitivity to low sucrose concentrations and that sucrose also promotes tillering independently of D14 and D3.

**Figure 6.**
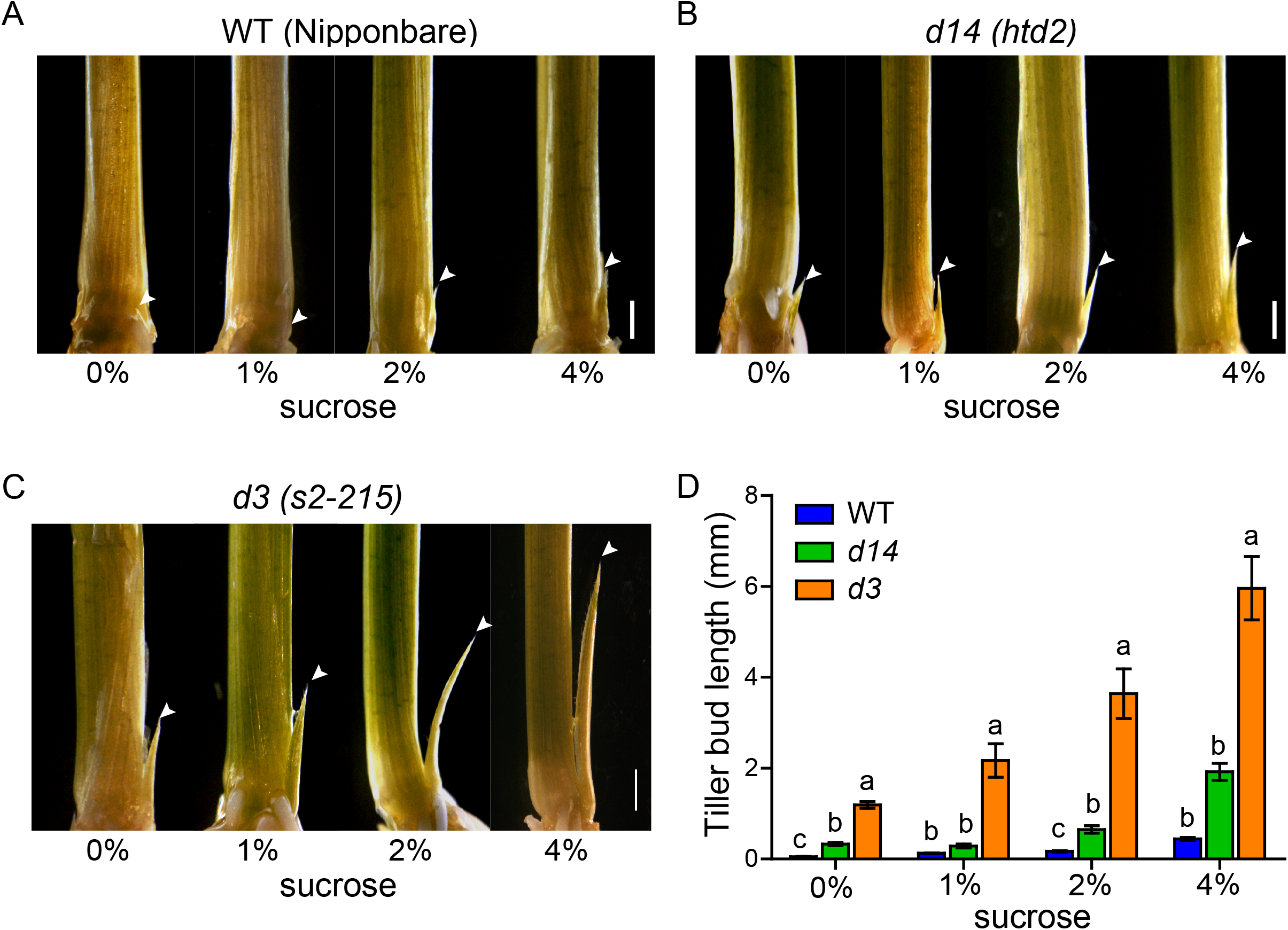
Growth of rice tiller buds in response to different sucrose concentrations. (**A**) Representative images of tiller buds of WT (Nipponbare), (**B**) SL signalling mutants *d14 (htd-2)* and (**C**) *d3 (S2-215)* grown on different sucrose concentrations for three weeks. (Scale bar= 1 mm). Arrowhead represents tiller bud used for measurement. (**D**) Tiller bud outgrowth (mm) in the WT (Nipponbare), *d14* (*htd-2*) and *d3* (*s2-215*) grown under the different sucrose concentrations for three weeks. Different lower-case letters denote significant differences, p<0.05, one-way ANOVA following Tukey’s test for multiple comparisons (compared for each sucrose concentration separately). Error bars represent ±SE (n > 8).

### *RMS4/PsD3/PsMAX2* is involved in the sucrose-induced bud outgrowth in garden pea

The role of sugars in bud release has been well described in garden pea, which is an established model eudicot for the study of shoot branching. In this species, decapitation triggers bud outgrowth through redistribution of sugars towards axillary buds, and sucrose feeding can trigger bud release (Mason *et al*., 2014; Fichtner *et al*., 2017). We explored whether a similar mechanism to what we observed in rice may occur in pea. To do so, we examined the expression of *PsD3 and PsD14* in pea (also known as *RMS4/PsMAX2* and *RMS3*, respectively) as well as their downstream target *PsBRC1*, the pea orthologue of *TB1*, in response to sucrose feeding and decapitation (Figure 7). Similar to the strongly sucrose-responsive expression of *D3* and *TB1* in rice, decapitation led to a decrease in *PsD3* and *PsBRC1* expression but not in that of *PsD14* (Figure 7A-C). Additionally, compared with sorbitol used as an osmotic control, sucrose feeding for 4 hrs through the petiole strongly inhibited *PsD3* expression (70%) and, to a lesser extent, *PsD14* expression (50%) (Figure 7D). *PsBRC1* expression was also repressed by sucrose but not significantly, compared with sorbitol at this time point (Figure 7D). The downregulation of expression of the sugar-repressible marker gene *PsDARK INDUCIBLE1* (*PsDIN1*) (Fujiki *et al*., 2001) indicates that the sucrose fed through the petiole reached the bud at 4 hrs. Altogether, these results support the hypothesis that *RMS4*/*PsD3*/*PsMAX2* is regulated by sucrose during bud outgrowth in pea, similar to that observed for *D3* in rice.

**Figure 7.**
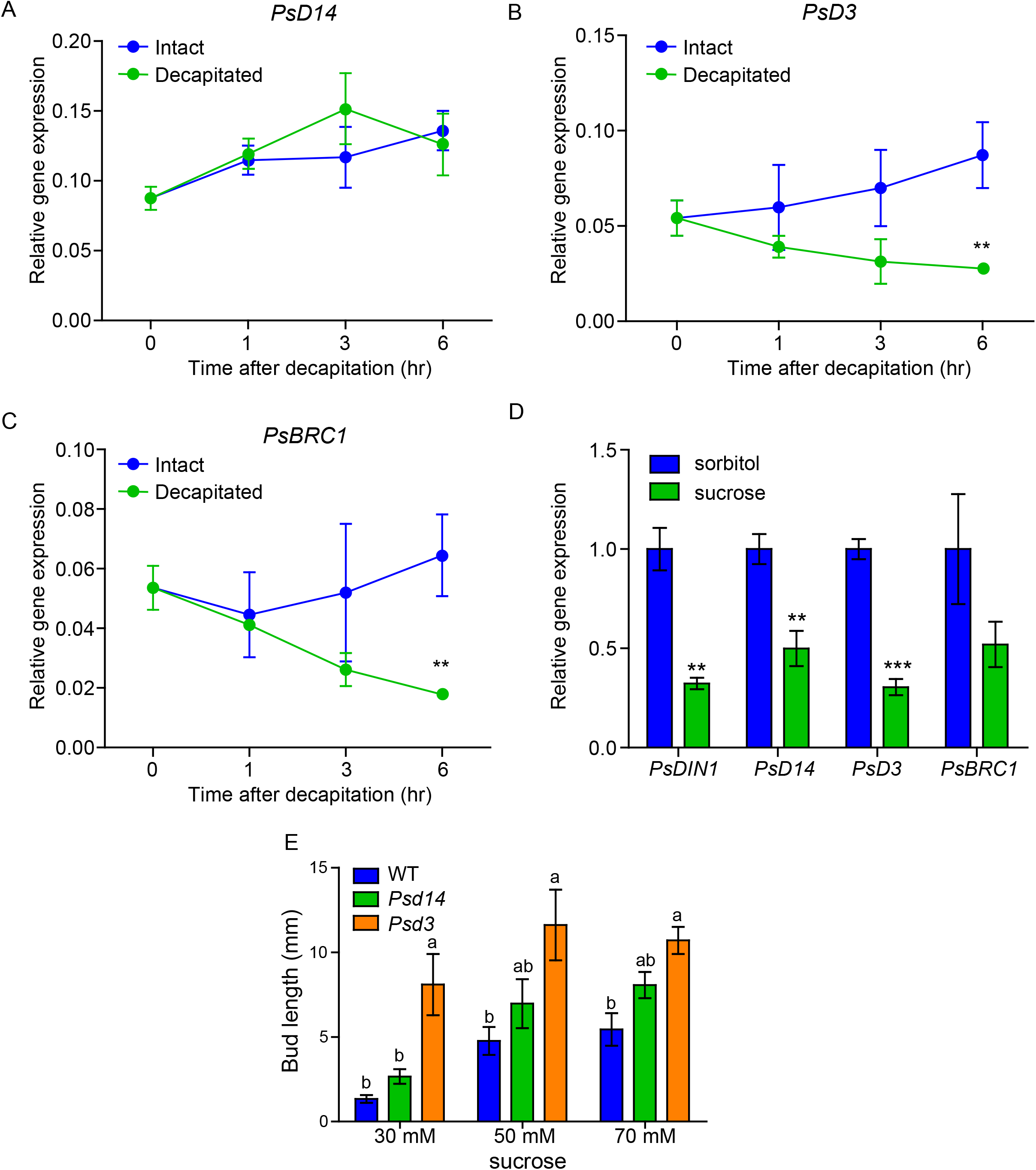
Effect of sucrose and decapitation on bud elongation and expression of SL pathway genes in pea buds. (**A**) Effect of decapitation on gene expression of *PsD14*, (**B**) *PsD3 (RMS4)* and (**C**) *BRC1* (*PsTB1*) at different time points in response to decapitation. Values are mean ± SE (n = 3 pools of 20 buds). (**D**) Expression of SL signalling genes in axillary buds fed with sucrose or sorbitol (osmotic control) through the petiole for 4h. Values are mean ± SE (n = 3 pools of 20 buds). Each replicate consists of 8 individual samples. Significant levels: ***p < 0.001; **p <0.01 indicated by Student’s *t*-Test. (**E**) Length of single-node pea buds of WT (Terese), *Psd14* (*rms3*) and *Psd3* (*rms4*) mutants grown *in vitro* with 30, 50 and 70 mM sucrose for 6 days. Different lower-case letters denote significant differences at each concentration (p<0.05, one-way ANOVA following Tukey’s test for multiple comparisons). Error bars represent ± SE (n = 8 individual buds).

We then tested responsiveness to sucrose of the *d14* and *d3* mutants in pea, also known as *rms3* and *rms4*, respectively. To achieve this, we grew pea single nodes on half-strength MS media supplemented with different sucrose concentrations (Figure 7E). Decreasing sucrose concentration from 50 mM to 30 mM inhibited bud elongation of the WT plants. As observed in rice, buds of the *d3* mutant were not as sensitive as the WT to lower sucrose concentrations (30 mM). However, contrary to what we observed in rice, buds of the *d14* mutant were not significantly different from the WT. Altogether these results indicate that disruption of *D3* leads to a lower bud response to decreased sucrose availability.

## Discussion

### Sucrose antagonises the inhibitory effect of strigolactones on tillering

Sucrose and SL play a crucial role in shaping plant architecture through their antagonistic action on bud outgrowth, as previously demonstrated in dicotyledonous plants like rose, pea and chrysanthemum (Dierck *et al*., 2016; Bertheloot *et al*., 2020). In the present study, we demonstrated that sucrose also promotes tillering and inhibits the impact of SL on this process in monocotyledonous plants. In rice, as in the previously mentioned species, the inhibitory effect of SL on bud outgrowth was almost totally prevented by high sucrose concentrations. The inhibitory effect of sucrose on SL perception is not limited to shoot branching as reported by recent studies showing that sucrose can also alleviate the effect of SL on dark-induced leaf senescence in rice (Takahashi *et al*., 2021) and bamboo (*Bambusa oldhamii*) (Tian *et al*., 2018).

The expression of the TCP transcription factor *BRC1* that inhibits shoot branching (Takeda *et al*., 2003; Aguilar-Martínez *et al*., 2007; Braun *et al*., 2012), has previously been reported to be repressed by sucrose in dicot species (Mason *et al*., 2014; Barbier *et al*., 2015b; González-Grandío *et al*., 2017; Otori *et al*., 2017; Wang *et al*., 2019b). Our observations have demonstrated that the expression of *TB1*, the *BRC1* homologue in monocots, is also repressed by sucrose in rice. Furthermore, D53 protein levels, which inhibit *TB1* gene expression, are increased by sucrose (Figure 2 A,D). Results in rice and *N. benthamiana* showed that SL-mediated D53 degradation was reduced by sucrose treatment (Figure 2 B,C; Supplementary Figure S5). This supports the hypothesis that the antagonistic effect of sucrose on SL-mediated bud inhibition (Dierck *et al*., 2016; Bertheloot *et al*., 2020) (Figure 1) is at least partly mediated through sucrose dampening SL-induced D53 degradation (Figure 8).

**Figure 8.**
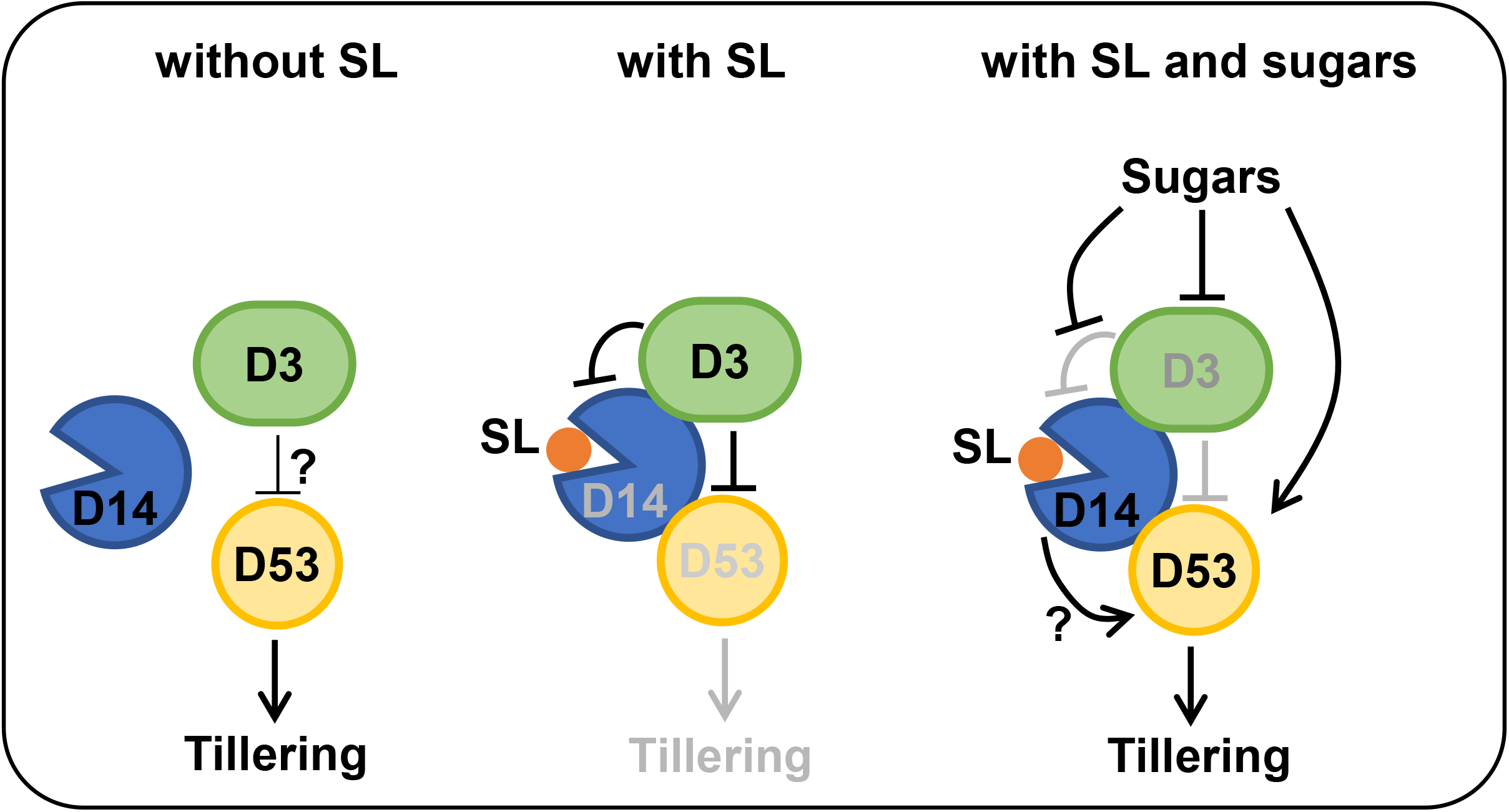
Proposed model of the interaction between sucrose and SL pathway. In absence of SL, D53 is not targeted by D14 and D3 and triggers tillering, although a residual SL-independent effect of D3 on D53 may exist. In presence of SL, a complex is formed between D14, D3 and D53, leading to the rapid degradation of D14 and D53, inhibiting tillering. In presence of SL and sugars, sugars repress *D3*, thus preventing the SL-induced D14 and D53 degradation and triggering tillering. Sugars also prevent the SL-induced D53 degradation through a D3-independent pathway and trigger a D14-dependent induction of D53 accumulation. The mechanism through which D14 promotes D53 accumulation remains unknown (hypotheses are given in the discussion).

### D3/MAX2 plays a key role in the antagonism between sugar availability and strigolactones

The F-box protein D3/RMS4/MAX2 plays an essential role in mediating the SL-dependent degradation of D53 protein through the SKP1–CULLIN–F-BOX (SCF) ubiquitin-proteasome system (Zhou *et al*., 2013; Zhao *et al*., 2014). Our data showed that over-expression of *D3*, but not of *D14*, prevented sucrose to antagonise the SL-induced D53 degradation and tillering inhibition (Figures 4 and 5), demonstrating the importance of D3 in modulating the tillering response to SL and sugar availability. In addition, the relative impact of sucrose on D53 accumulation in presence of SL was much stronger in the WT than in the *d3* mutant (Figure 4I), supporting this conclusion. As previously observed in sorghum, rose and arabidopsis (Kebrom *et al*., 2010; Barbier *et al*., 2015b, 2021), sugar availability suppresses *D3/MAX2* gene expression in rice and pea and this effect is stronger and more consistent than for *D14* (Figure 3C, 3D and 7D). In addition, we did not observe a direct effect of sucrose on D3 protein levels (Figure 3F), showing that sucrose preferentially regulates *D3* transcription. However, sucrose might also act through D3 protein by regulating the switch between the two conformational states of this protein which has been reported to modulate the binding affinity between D3 and D14 (Shabek *et al*., 2018).

Some evidence suggest that D3/MAX2 may retain a function independent of SL. In field conditions, the *d3 (s2-215)* mutant (Patil *et al*., 2019) showed a more severe dwarf and high tillering phenotype compared with the *d14 (htd-2)* mutant (Liu *et al*., 2009), both being loss-of-function mutants developed from the same background (Nipponbare) (Supplementary figure S7). In both rice and pea buds, we could also observe this difference between *d14* and *d3* mutants, particularly under low sugar availability (Figure 6 and 7). This stronger impact of *D3* mutation compared with *D14* mutation or SL-deficiency has been reported in different species and in different developmental processes (Umehara *et al*., 2008; Hayward *et al*., 2009), showing that our observations are not due to a specific allele, and the difference between *D3* and *D14* mutations is conserved in diverse species. It was previously reported in arabidopsis, that over-expression of *MAX2* could partially suppress decapitation-induced branching in a SL-deficient background (Stirnberg *et al*., 2007), supporting the hypothesis that D3/MAX2 may retain a function independently of SL.

Besides mediating SL signalling, D3/MAX2 has been shown to mediate the impact of karrikins in different developmental processes, including seed germination and root development (Nelson *et al*., 2011; Waters *et al*., 2012). However, karrikins have been reported to have no effect on shoot branching (Nelson *et al*., 2011). It is therefore unlikely that the SL-independent effect of D3/MAX2 on branching is dependent on karrikin signalling. Interestingly, MAX2 was recently reported to be involved in CO2 signalling in arabidopsis (Kalliola *et al*., 2020). In addition, new proteins interacting with MAX2 were recently discovered (Struk *et al*., 2021), showing that the regulation of MAX2 signalling is more complex than previously thought. Altogether, these observations have demonstrated that D3/MAX2 is an important regulator in the antagonistic regulation of shoot branching and tillering by sugars and strigolactones (Figure 8).

The molecular mechanism through which sugars inhibit *D3/MAX2* expression is unknown. However, recent studies have highlighted that sugars regulate shoot branching through different sugar-signalling components such as Tre6P (Fichtner *et al*., 2017, 2021a) and HXK1 (Barbier *et al*., 2021). Interestingly, it was shown that *MAX2* expression negatively correlates with *HXK1* expression and with sugar availability in arabidopsis (Barbier *et al*., 2021). Moreover, HXK1 deficiency leads to an upregulation of *MAX2* expression (Barbier *et al*., 2021), suggesting that the HXK1-signalling pathways may be involved in sugar regulation of *MAX2* expression. However, HXK1 signalling is specific to glucose, and our previous data suggests that a sucrose-specific signalling pathway was also involved in shoot branching (Barbier *et al*., 2015b). Since Tre6P has been shown to be a sucrose-specific signal (Fichtner & Lunn, 2021), it is reasonable to assume that this signalling component is also involved in the regulation of *MAX2* expression by sugars. In addition, other sugar signalling pathways have been shown to regulate *BRC1* expression via transcriptional (Wang *et al*., 2021) and post-transcriptional mechanisms (Wang *et al*., 2019b). It is therefore possible that sugar-signalling mechanisms independent of HXK1 and Tre6P may also be involved in the regulation of *MAX2* expression. Future studies need to be done to understand how sugars regulate *MAX2* expression at the molecular level.

### Sucrose prevents SL-induced D53 and D14 degradation to promote tillering

The transcriptional inhibition of *D3/MAX2* by sucrose in diverse plants suggests that the effect of sucrose on D53 may be mediated at least partly via *D3*. This hypothesis is strongly supported by the evidence that sucrose is able to alleviate GR24-mediated D53 degradation in a *D14* over-expressing line, but not in a *D3* over-expressing line (Figure 4 D, F). The inability of sucrose to alleviate LUC-D53 protein degradation when co-transfected with the *35Sp:OsD3* construct provides further supports that sucrose acts, at least partly, through D3 to regulate D53 (Figures 4H and 8; Supplementary Figure S5).

In addition to degradation of D53 protein, D3 is also responsible for SL-mediated degradation of D14 protein in rice (Hu *et al*., 2017). We thus measured the effect of sucrose on HA-tagged D14 protein levels. In the absence of sucrose, GR24 completely degraded the D14 protein within 12 hrs, whereas in the presence of 4% sucrose, GR24 failed to degrade the D14 protein (Figure 3E). This result is important for two reasons. Firstly, it shows that the effect of sucrose on D53 protein levels could not be attributed to a negative effect of sucrose on D14 protein levels. Secondly, it suggests that the effect of sucrose on D53 is likely to be mediated by D3 since protein accumulation pattern of D14 and D53 reflects what would be expected if *D3* was down-regulated (Hu *et al*., 2017) (Figure 8). Contrary to the HA-D14 protein, sucrose did not show any positive or negative effect on HA-D3 protein levels, indicating sucrose acting through transcriptional regulation of *D3* rather than through regulation of D3 protein stability (Figure 3F).

Interestingly, we observed an over-accumulation of D53 protein in the *D14* over-expression line (Figure 4A-D, Supplementary Figure S8), which further led to increased sucrose-induced tillering in this line (Figure 5A-B). Considering the dual function of the D3 protein in modulating both SL-induced D53 and D14 protein degradation (Chevalier *et al*., 2014; Soundappan *et al*., 2015; Wang *et al*., 2015; Hu *et al*., 2017), it can be hypothesised that there may be a competitive effect between D53 and D14 in binding to the D3 protein. Elevated D14 protein levels in the *D14* over-expressing line would recruit more D3 protein, creating a deficit for D53 protein degradation. Whether and tillering remains to be determined. Interestingly, in pea, D14 was reported to be a mobile protein between root and shoot (Kameoka *et al*., 2016). In addition, D14 was reported to be highly abundant in the phloem in rice (Aki *et al*., 2008) and arabidopsis (Batailler *et al*., 2012). Combined with our data, this suggests that the role of D14 in SL signalling is more complex than previously thought. More work is needed to test whether sucrose acts directly on D14 to increase its protein levels and increase D53 levels and to better understand the role of D14 in SL signalling.

### Sucrose also promotes tillering through a D3-independent pathway

Our results also show that sucrose can increase D53 protein levels (Figure 4J) and inhibit *TB1* expression (Supplementary Figure S6) in the *d3* background, without addition of exogenous SL. In addition, bud elongation was also stimulated by sucrose in the *d3* background (Figure 6I-J), showing that at least one other pathway, independent of D3, is involved in the sucrose-induced tillering. One straightforward explanation could be that the trophic properties of sucrose also play a role in inducing D53 accumulation and tillering. However, other hypotheses could also explain these observations. SL were reported to lead to the degradation of D14 by the same process as that leading to D53 degradation (Hu *et al*., 2017). Our results also showed that sucrose prevents SL from degrading D14 (Figure 3E), which makes senses if sucrose prevents the degradation of D53 by inhibiting *D3* (see section above). This could also be explained by sucrose inhibiting the proteasome complex required for D3-dependent D14 and D53 degradation. We also showed that increasing D14 led to higher levels of D53 (Figure 4 A,D and Supplementary Figure S8). It could therefore be possible that sucrose prevents the SL-induced D14 degradation to promote D53 accumulation (Figure 8). This could be achieved by inhibiting the binding of SL to D14 or by preventing D3 from binding to D14. Recent findings in pea suggested that CK promotes PsSMXL7 protein accumulation, the homologue of the rice D53, through increasing the expression of *PsSMXL7* gene (Kerr *et al*., 2021). Since sucrose was reported to promote CK synthesis in different species (Barbier *et al*., 2015b; Kiba *et al*., 2019; Salam *et al*., 2021), it would be tempting to hypothesise that sucrose induces D53 protein levels through a CK-mediated increase of *D53* expression. However, in our conditions, sucrose did not promote *D53* expression, and even decreased its transcription (Figure 3B), excluding this hypothesis.

Whether sucrose acts through D53 to regulate *TB1* expression independently of D3 remains to be determined (Figure 3A and Supplementary Figure S6). As mentioned above, sugar availability was reported to induce CK accumulation (Barbier *et al*., 2015b; Kiba *et al*., 2019; Salam *et al*., 2021), and CK were reported to inhibit *BRC1* expression (Dun *et al*., 2012; Roman *et al*., 2016), which could explain our observations. It was also reported in rose calli, that sucrose regulates *TB1/BRC1* expression through specific elements on the 3’UTR of this gene (Wang *et al*., 2019b), showing that sugars interact with SL signalling through multiple pathways. More studies are needed to understand the complete mechanism underpinning sugar-promoted tillering/branching and the interactions between SL and sugars.

### Conclusion

The present study demonstrates that sugar availability and strigolactone signalling interact during the control of shoot branching, and that D3 plays a role in this interaction but that other pathways independent of D3 may also exist. In addition, our study shows that the protein levels of D14 and D3 impact tillering in opposite ways, which should prompt further studies to investigate the regulation of the components in the SL regulation of plant development. SL and sugars are systemic signals and their levels in plants are tightly regulated by environmental cues such as light, moisture and nutrient status (Yoneyama *et al*., 2007; Yoneyama *et al*., 2012; Lemoine *et al*., 2013; Kapulnik & Koltai, 2014). Their interactions during the control of bud outgrowth therefore represents an important regulatory node in the control of plant architecture in response to the environment. Our study therefore provides an interesting opportunity to manipulate tillering and ultimately improve crop management and yields through molecular engineering and crop selection. Nonetheless, more work is needed to fully understand the mechanism(s) through which sucrose interact with D14, D3 and D53, and to better understand the unexpected promoting role of D14 in tillering.

## Acknowledgments

We would like to thank Catherine Rameau, Alexandre de Saint-Germain and Steven Smith for fruitful discussions about the manuscript. Work carried out in China was supported by grants from National Natural Science Foundation of China (31670279, 31271311 to Xueyong Li) and the Ministry of Agriculture of China (2016ZX08009-003 to Xueyong Li). SBP is a PhD scholar supported by the China Scholarship Council (CSC) & Ministry of Human Resource Development, Government of India. The work carried out in Australia was funded as part of a Researcher Development initiative and by the Australian Research Council (Georgina Sweet Laureate Fellowship FL180100139 to Prof. C. A. Beveridge).

## Supplementary Material

**Supplementary Table S1.**
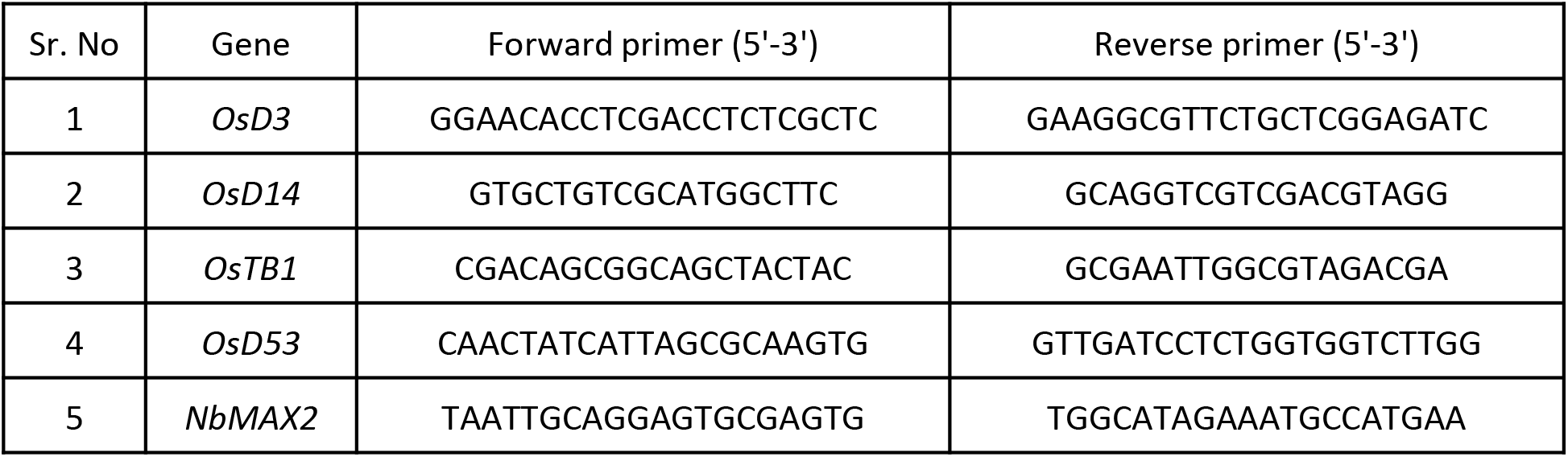
Primers used in RT-qPCR analysis

**Supplementary Figure S1.**
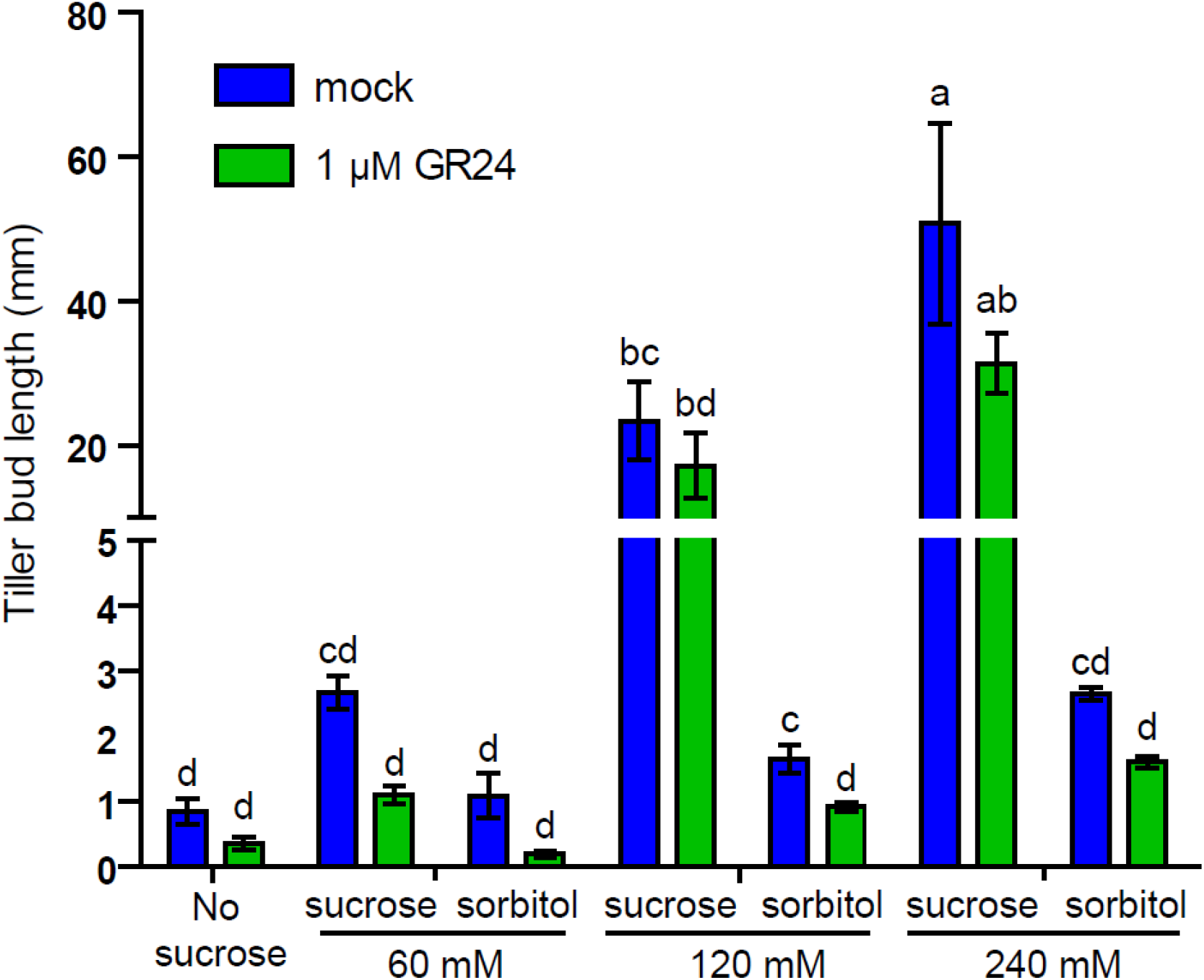
Effect of sucrose and sorbitol on tillering. Length of tiller buds from rice plants fed hydroponically with different sucrose and sorbitol concentrations [No sucrose, 2% (∼60 mM), 4% (∼120 mM) and 8% (∼240 mM)] with or without 1μM GR24. Different lower case letters denote significant differences, p<0.05, one-way ANOVA following Tukey’s test for multiple comparisons. Error bars represent ± SE (n > 8).

**Supplementary Figure S2.**
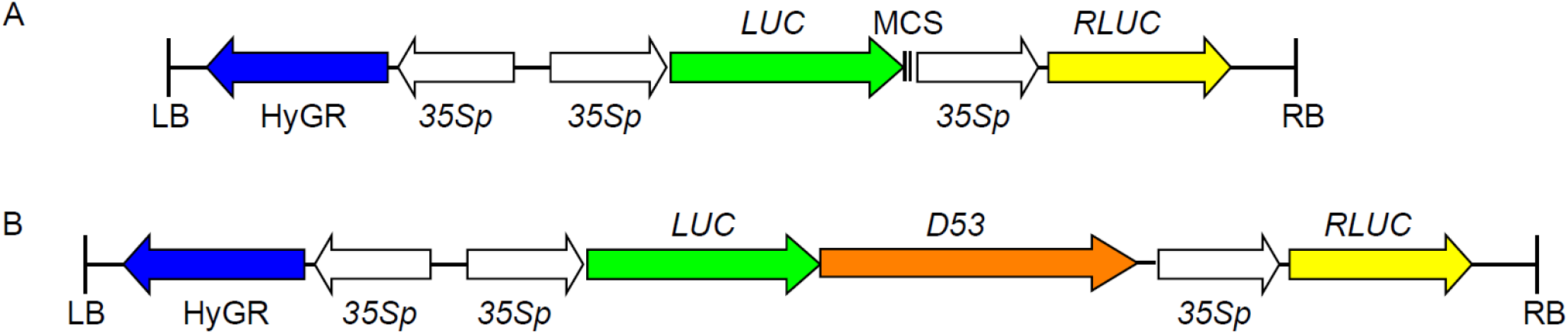
Map of the vector used for the D53 protein degradation experiment performed in tobacco leaves. (**A**) Original vector (pCAMBIA 1200-R-LUC) consisting of the Fire fly luciferase and Renilla luciferase sequence driven by the CaMV 35S promoter. (**B**) The vector with the D53 protein fused at the C-terminal end of the Fire fly luciferase coding sequence driven by the CaMV 35S promoter in addition to the Renilla luciferase gene driven by the CaMV 35S promoter.

**Supplementary Figure S3.**
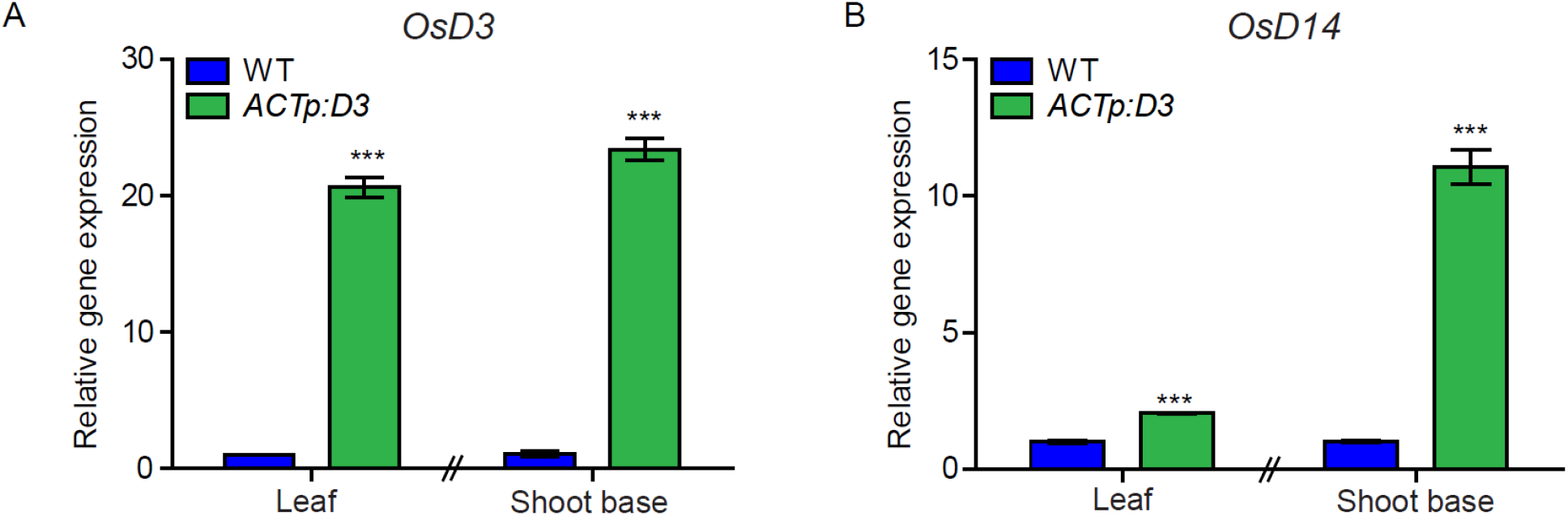
Confirmation of gene expression of *OsD3* and *OsD14* in the corresponding overexpressing transgenic lines. Two-week-old rice seedlings were used to detect the expression of *OsD3* in the WT (GSOR300192), and *ACTp:D3* (*D3* over-expression line B11-15) and *OsD14* expression in the WT (GSOR300192) and *ACTp:D14* (*D14* over expression line B8-14). Significant levels (Compared to the WT), ***p <0.001 indicated by Student’s t-Test.

**Supplementary Figure S4.**
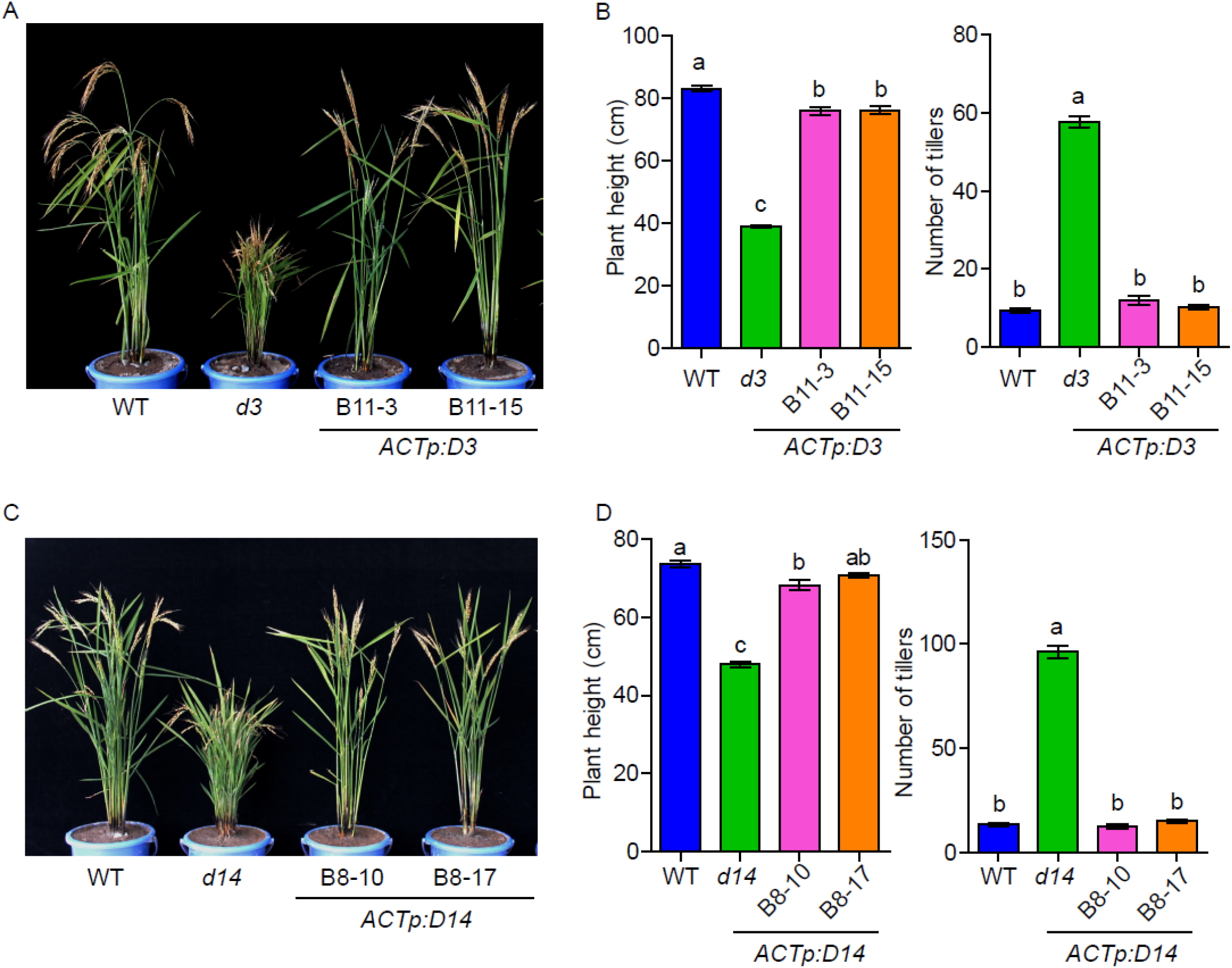
Phenotypes of the d3 and d14 loss-of-function mutants in comparison with WT in the field condition.. (**A**) Phenotypes of WT (GSOR300192), *d3 (gsor300097)* and *ACTp:D3* transgenic plants at maturity stage in field condition. (**B**) Plant height (cm) and tiller numbers of the Phenotypes of WT (GSOR300192), *d3 (gsor300097)* and *ACTp:D3* transgenic lines at maturity stage in the field condition. Values are mean ±SE (n=20). (**C**) Phenotypes of WT (GSOR300192), *d14 (gsor300183)* and *ACTp:D14* transgenic plants at maturity stage in field condition. (**D**) Plant height (cm) and tiller numbers of the WT (GSOR300192), *d14 (gsor300183)* and *ACTp:D14* transgenic lines at maturity stage in the field condition. Different lower case letters denote significant differences (p<0.05, one-way ANOVA following Tukey’s test for multiple comparisons). Error bars represent ± SE (n=20).

**Supplementary Figure S5.**
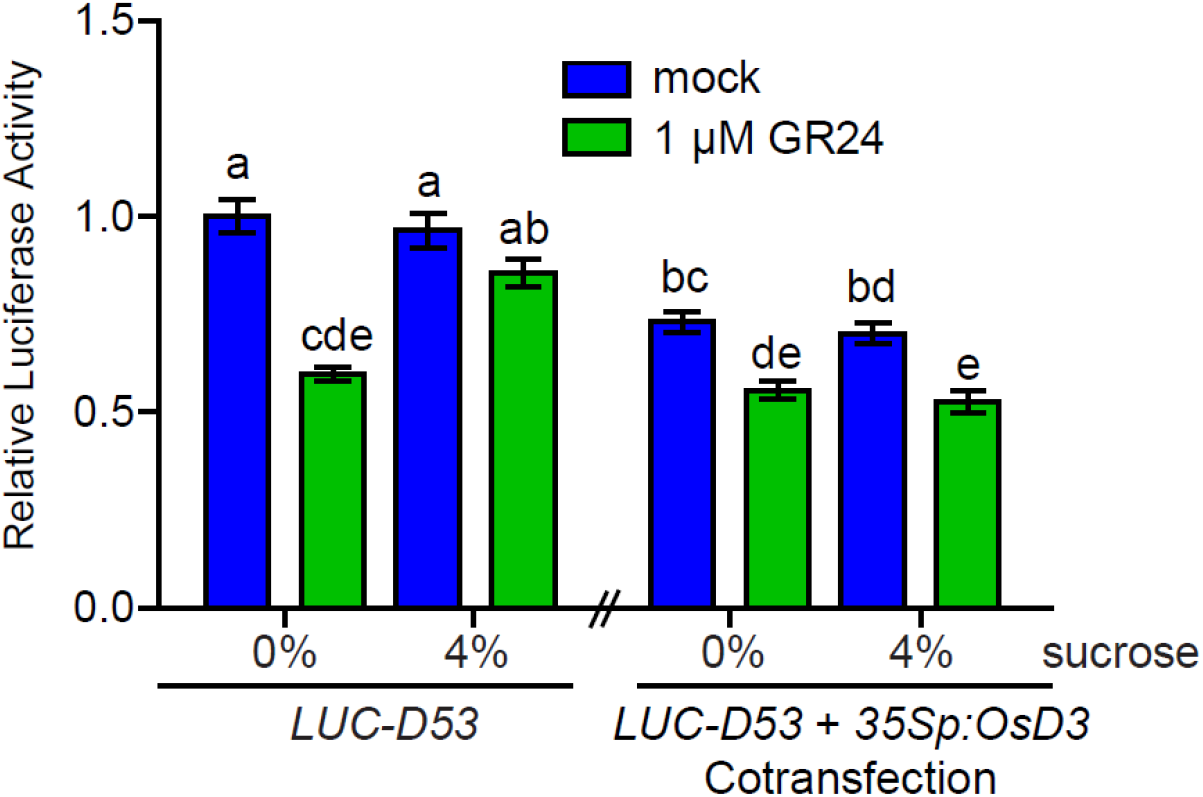
Effect of sucrose and GR24 on transiently expressed luciferase-D53 protein alone and co-transfected with 35S-OsD3 protein in *Nicotiana benthamiana* leaves. Luciferase readings were normalised with renilla luciferase readings. Different lower-case letters denote significant differences (p<0.05, one-way ANOVA following Tukey’s test for multiple comparisons). Error bars represent ± SE (n=6).

**Supplementary Figure S6.**
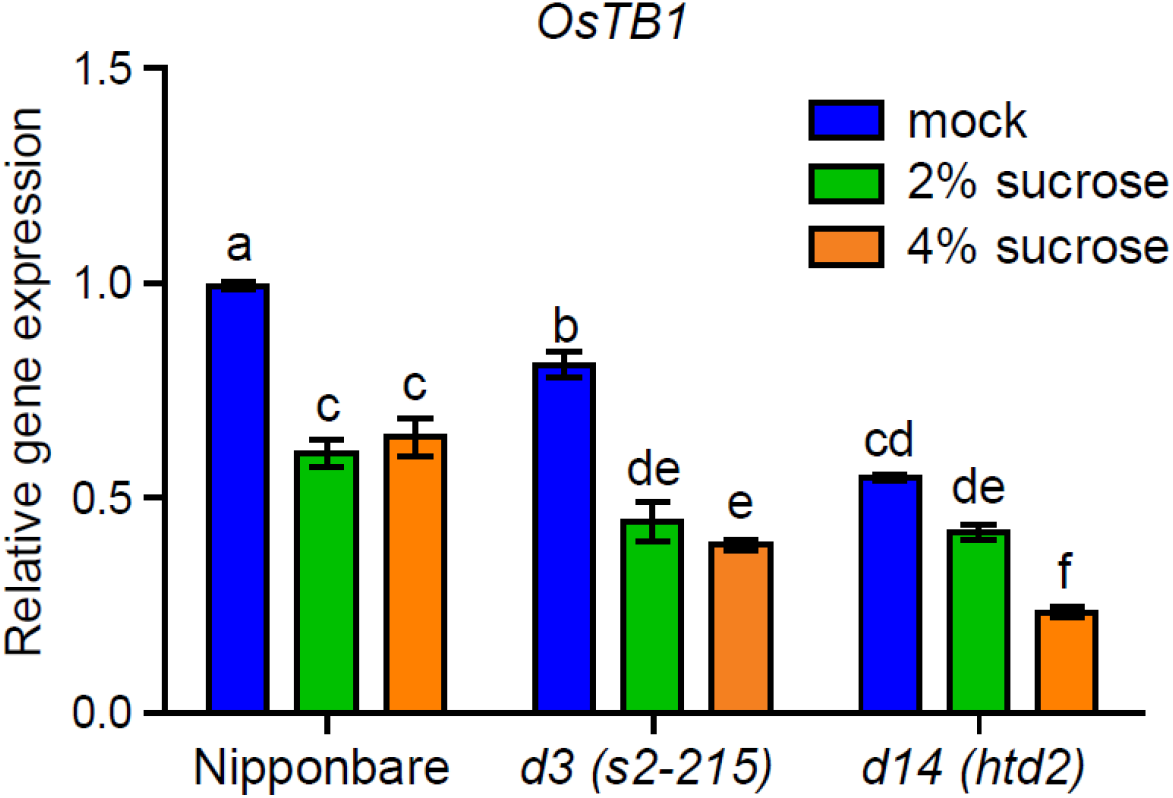
*OsTB1* expression in the rice calli treated with different concentrations of sucrose for 24 hrs. Different lower-case letters denote significant differences (p<0.05, one-way ANOVA following Tukey’s test for multiple comparisons). Error bars represent ± SE (n=6)).

**Supplementary Figure S7.**
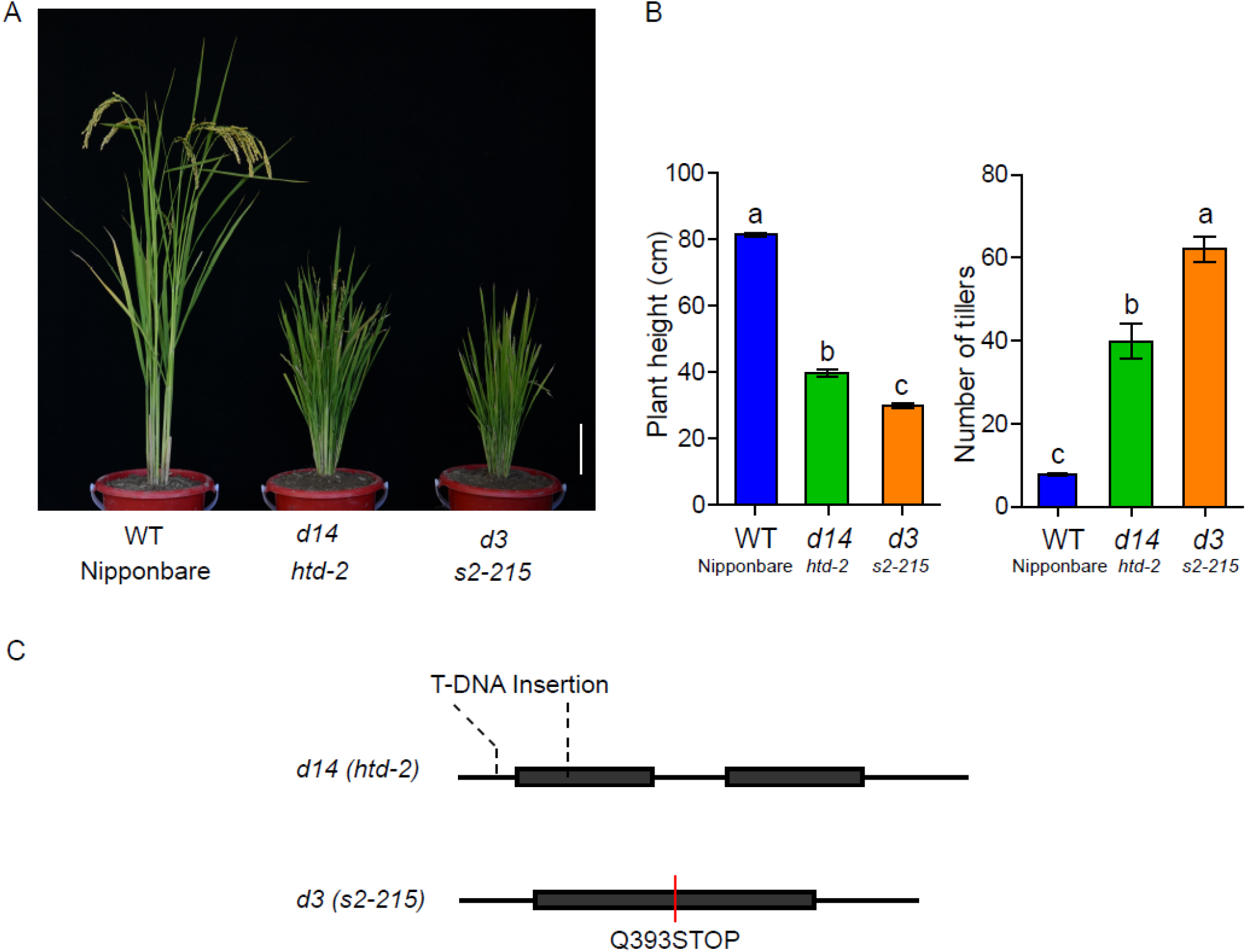
Phenotypes of the *d3* and *d14* loss-of-function mutants in comparison with WT in the field condition. (**A**) Phenotypes of WT (Nipponbare), *d14 (htd-2)* and *d3 (s2-215)* mutants grown in the field condition. Scale bar=5 cm. (**B**) Plant height (cm) and tiller number of WT, *d14* and *d3* grown in the field condition. Different lower case letters denote significant differences (p<0.05, one-way ANOVA following Tukey’s test for multiple comparisons). Error bars represent ± SE (n =8). (**C**) Mutation points of *d14 (htd2)* and *d3 (s2-215)* in the gene structure (black boxes represent exon and black line represent intron).

**Supplementary Figure S8.**
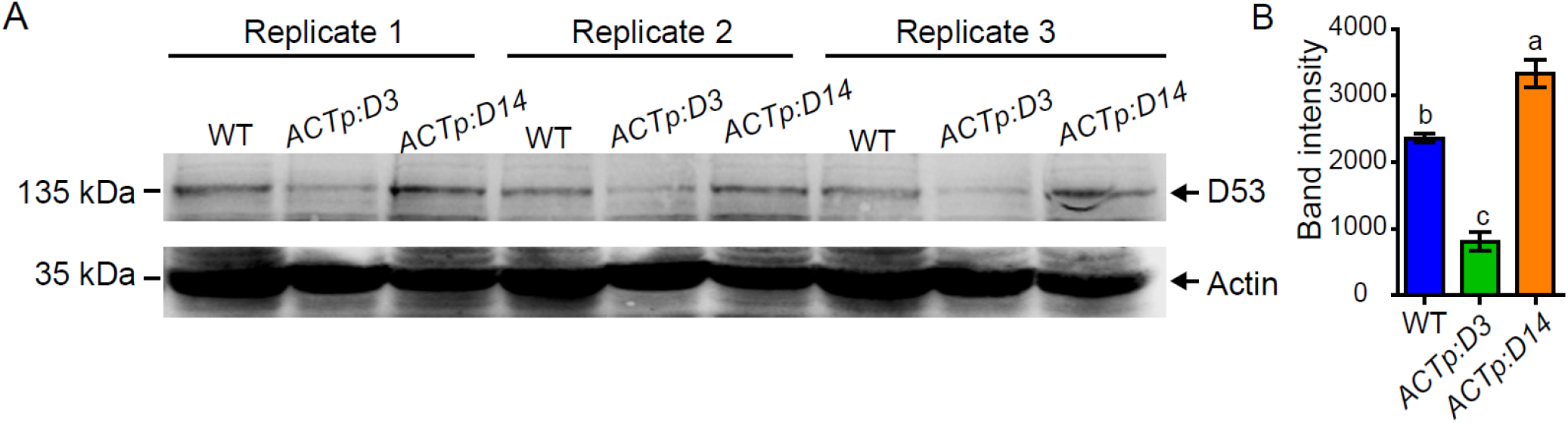
Comparative analysis of D53 protein levels in different backgrounds. (**A**) D53 protein levels in WT, *ACTp:D3* and *ACTp:D14* over-expressing lines grown on 4% sucrose plates detected by immunoblotting with an anti-D53 polyclonal antibody. (**B**) Average band intensity of the D53 proteins quantified in the WT, *ACTp:D3* and *ACTp:D14* over-expressing lines using Image J software. Values are mean ± SE (n=3), Different lower case letters denote significant differences, p<0.05, one-way ANOVA following Tukey’s test for multiple comparisons.

## Notes

### Competing Interest Statement

The authors have declared no competing interest.

### Summary of Updates

Shoot branching, a major component of shoot architecture, is regulated by multiple signals. Previous studies have indicated that sucrose may promote branching through suppressing the inhibitory effect of the hormone strigolactone (SL). However, the molecular mechanisms underlying this effect are unknown. Here we used molecular and genetic tools to identify the molecular targets underlying the antagonistic interaction between sucrose and SL. We showed that sucrose antagonises the suppressive action of SL on tillering in rice and on the degradation of D53, a major target of SL signalling. Sucrose inhibits the expression of D3, the orthologue of the arabidopsis F-box protein MAX2 required for SL signalling. Over-expression of D3 prevents sucrose from inhibiting D53 degradation and enabled the SL inhibition of tillering under high sucrose. Sucrose also prevents SL-induced degradation of D14, the SL receptor involved in D53 degradation. Interestingly, D14 over-expression enhances D53 protein levels and sucrose-induced tillering. Our results show that sucrose inhibits SL perception by targeting key components of SL signalling and, together with previous studies reporting the inhibition of SL synthesis by nitrate and phosphate, demonstrate the central role played by strigolactones in the regulation of plant architecture by nutrients.

